# MAASTY: A (dis)ordered copolymer for structural determination of human membrane proteins in native nanodiscs

**DOI:** 10.1101/2024.08.19.608676

**Authors:** Ciara F. Pugh, Lukas P. Feilen, Dušan Zivković, Kaia F. Præstegaard, Charalampos Sideris, Neil J. Borthwick, Casper de Lichtenberg, Jani R. Bolla, Anton A. A. Autzen, Henriette E. Autzen

## Abstract

Amphiphilic copolymers capable of extracting membrane proteins directly from cellular membranes into ”native nanodiscs” offer a simplified approach for preparing membrane proteins in lipid nanodiscs compared to approaches that rely on detergent. Amphiphilicity, length, and composition influence the performance of copolymers, in addition to the protein itself and the purification conditions used. Here, we report a copolymer composed of methacrylic acid and styrene, which we term MAASTY, leveraging the inherent monomer reactivity ratios to create an anionic copolymer with a statistical distribution of monomers. We show that MAASTY can be used for high-resolution structural determination of a human membrane protein with single particle cryo-electron microscopy, preserving endogenous lipids including cholesterol and exhibiting an enrichment of phosphatidylinositol. Moreover, MAASTY copolymers effectively solubilize a broad range of lipid species and a wide range of different, eukaryotic membrane proteins from mammalian cells. We find that MAASTY copolymers are promising as effective solubilizers of membrane proteins and offer a chemical platform for structural and functional characterization of membrane proteins in native nanodiscs.

## Introduction

Integral membrane proteins represent an important class of therapeutic targets necessitating comprehensive understanding of their functional properties and structural attributes to facilitate rational drug development^1^. Specifically, structural characterization of integral membrane proteins has witnessed substantial progress the past decade, particularly driven by the technological advancements in single particle cryoelectron microscopy (cryo-EM)^2^. High-resolution cryo-EM has sparked an increased focus on reconstituting integral membrane proteins in lipid nanodiscs to effectively mimic native-like membrane environments^3^. The dominating approach for incorporating membrane proteins into lipid nanodiscs relies on detergent solubilization followed by subsequent re-lipidation, enabling the insertion of the membrane protein into a synthetic lipid bilayer encased by membrane scaffold proteins (MSPs) or saposin (Fig. 1a, black arrows)^4,5^. While reconstitution into synthetic nanodiscs facilitates the study of membrane proteins in a lipid environment, it inevitably comes at a cost of the native lipids, which are either partially or completely substituted by synthetic lipids. Consequently, only native lipids that exhibit exceptionally high affinity for the target protein remain associated with the reconstituted membrane protein^6–8^. Importantly, these reconstitution methods for the use of synthetic nanodisc systems disrupt the inherent asymmetry of the native membrane, may destabilize the membrane protein of interest through use of detergent-based extraction, and altogether, necessitate extensive and time-consuming optimization.

**Fig. 1:**
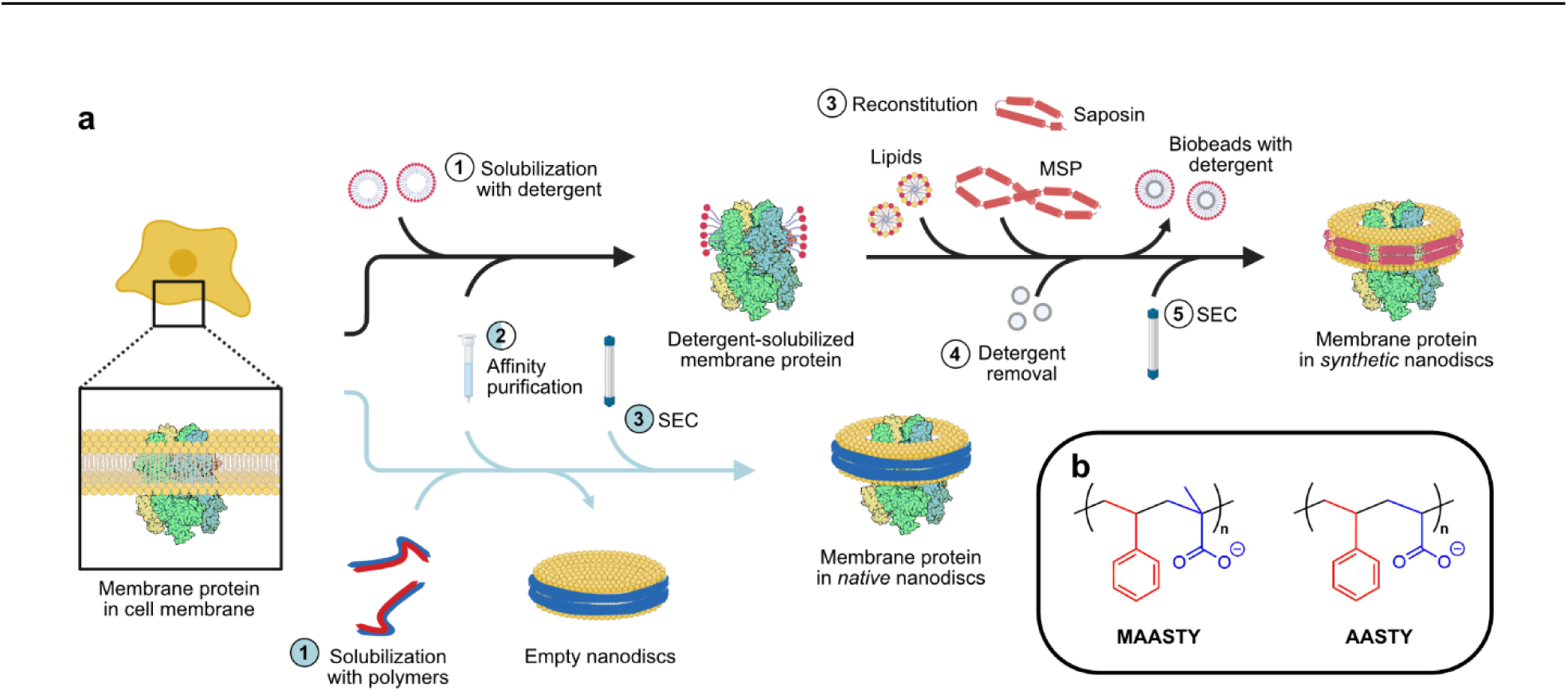
Overview of membrane mimetics systems and chemical composition of MAASTY copolymers. **a** Overview of the synthetic (white circles, black arrows) and native nanodisc (blue circles, blue arrows) approaches. Reconstitution into synthetic nanodiscs is typically carried out using detergents (steps 1–5, white circles). First, cells are treated with detergent to solubilize the proteins (step 1). The detergent-solubilized, recombinant protein is then purified using affinity chromatography (step 2), then reconstituted into a lipid bilayer by adding detergent-solubilized lipids and MSP or saposin (step 3) during which detergent is removed with bio-beads (step 4). Finally, aggregates and/or empty discs are removed by size-exclusion chromatography (SEC, step 5). In the native nanodisc approach, membrane proteins are purified with amphiphilic copolymers, leaving the membrane protein in a nanodisc that contains endogenous or native lipids (steps 1–3, blue circles). First, cells are incubated with the copolymer (step 1), during which the native nanodiscs form. Next, the protein in native nanodiscs is purified by affinity chromatography (step 2) and SEC (step 3). **b** Chemical structures of MAASTY characterized in this study and AASTY from previous studies^18,38^. MAASTY is composed of MAA (blue) and STY (red), while AASTY is composed by AA (blue) and STY (red).

Nanodisc-forming copolymers, characterized by their amphipathic nature, have emerged as a promising alternative to traditional approaches for reconstituting integral membrane proteins into lipid nanodiscs^9–14^. These polymers are able to dissolve membranes and directly embed integral membrane proteins into what are coined ’native nanodiscs’, bypassing the requirement for initial detergent extraction before lipid reconstitution (Fig. 1a, blue arrows). Although the native nanodisc method was established nearly two decades ago^12^, the method remains in an early stage of development; nevertheless, it has demonstrated significant potential for cryo-EM studies and functional characterizations of various membrane proteins expressed in both bacterial^9,15–17^ and mammalian cells^18–22^. However, despite the promise of the native nanodisc method, the limited number of high-resolution structures of membrane proteins embedded in native nanodiscs indicates certain limitations inherent to the catalog of currently available copolymers. The initial adoption of poly(styrene-co-maleic acid) (SMA) copolymers marked a significant milestone in the development of the native nanodisc technology, and their continued utilization remains extensive^12,23–25^. However, SMA suffers from notable drawbacks, such as its negative charge and high dispersity in molecular weight, limitations that collectively restrict its applicability in structural and functional characterization of membrane proteins^26–28^. Alternative copolymers aiming to address these challenges include poly(diisobutyleneco-maleic acid) (DIBMA)^29^, polymethacrylate (PMA)^30^, stilbene-maleic anhydride (STMA)^31^, pentyl-inulin^32^, poly(acrylic acid-co-styrene) (AASTY)^18^, in addition to various SMA derivative copolymers to mention a few^33–36^.

Here, we explore the use of a copolymer for native nanodisc formation, poly(methacrylic acid-co-styrene) composed of methacrylic acid (MAA) and styrene (STY), which we term MAASTY, and its applicability toward forming native nanodiscs. We hypothesized that MAASTY copolymers act as effective nanodisc-forming polymers on account of their structural similarity to SMA and AASTY and their highly alternating behavior due to the reactivity ratios between MAA and STY in radical polymerizations.

MAASTY was developed on the framework of the copolymer AASTY^18^, composed of acrylic acid (AA) and STY, but differing by a methyl on the backbone of the acid bearing monomer (Fig. 1b). This methyl influences the monomer reactivity ratios and the linear sequence of monomers in the resulting copolymers, as well as their overall amphiphilicity due to the hydrophobicity of the methyl group. Similar to AASTY, MAASTY can be synthesized by reversible addition-fragmentation chain-transfer (RAFT) polymerization, allowing control of the copolymer length and the dispersity of size compared to free-radical polymerization^37^. We decided to investigate MAASTY because of its moderately irregular monomer composition and reduced composition drift in bulk polymerizations, enabling the exploration of a broader range of homogeneous monomer compositions compared to AASTY.

We demonstrate that MAASTY can be used to extract and purify the human transient potential melastatin 4 (hTRPM4) ion channel from mammalian cells, allowing determination of its high-resolution structure with single particle cryo-EM. Using fluorescent size-exclusion chromatography (FSEC), we show that MAASTY copolymers extract a wide range of eukaryotic membrane proteins composed of different numbers of subunits in native nanodiscs and form nanodiscs independent of the bulk lipid head group charge, thus expanding the application potential of MAASTY copolymers across a variety of systems with intricate membrane lipid compositions.

## Results

### Synthesis and chemical characterization of MAASTY

Motivated by our previous work on AASTY copolymers^18,38^, we synthesized six MAASTY copolymers with a broad range of polymer compositions, varying the initial MAA monomer composition in the reactions from 40-65% whilst aiming for a molecular weight between 7 and 8 kDa (Table 1). The MAASTY copolymers are named according to their M_n_ (number-average molecular weight) and initial MAA fraction. In radical polymerizations, a given monomer composition in a copolymerization will not necessarily result in copolymers with identical polymer compositions due to relative differences in the monomer reactivity ratios. This is evident from comparing the final MAA fraction (MAA frac) in the polymers (Table 1) with the initial MAA fraction. Monomers are consumed at different rates, leading to a composition drift of monomers along the chain. The composition drifts are simulated in Fig. 2a for libraries of MAASTY and AASTY copolymers using their respective reactivity ratios (MAASTY; MAA: 0.4065, STY: 0.2911, AASTY; AA: 0.21, STY: 0.082)^39–41^. The copolymer composition drifts are displayed for initial monomer compositions (40-65%) of MAA or AA. As shown in Fig. 2a, AASTY initially has similar copolymer compositions across the simulated starting conditions due to its highly alternating nature, resulting in a significant drift in polymer composition at higher conversions. The compositions of MAASTY, on the other hand, are more dispersed at all conversions. This moderately alternating nature of MAASTY is further highlighted by its determined simulated chemical correlation factor, *λ*, of approximately −0.5 for all tested copolymers (Table 1). This behavior is also apparent when comparing the regularity of monomer sequences as run lengths of each monomer (Fig. 2b). Here, a perfectly alternating copolymer would have a run length of 1, with any deviation indicating a higher degree of randomness. With a run length of 1.29, AASTY is highly alternating in terms of the monomer sequence, while MAASTY is more irregular with a run length of 1.68. A similar pattern can be observed in the product of the reactivity ratios (*r*_1_*r*_2_) for both copolymers, as *r*_1_*r*_2_ for AASTY is 0.017^41^, whereas the *r*_1_*r*_2_ for MAASTY is 0.119^39^. Ultimately, this moderately alternating nature of the MAASTY co-polymerization allows for synthesis of a broader range of compositions without a significant drift in comparison to AASTY copolymerization.

**Fig. 2:**
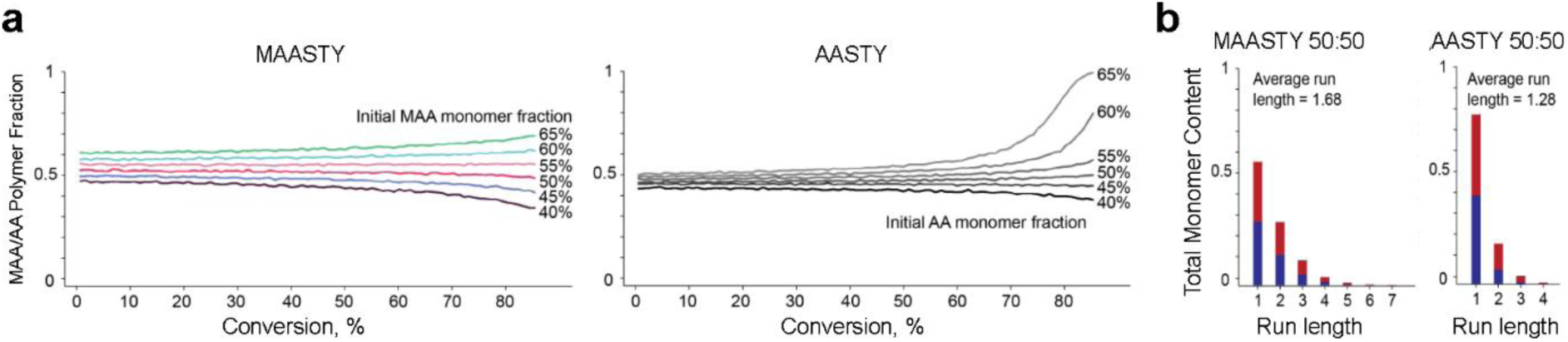
Simulated composition drift and run length analyses of MAASTY and AASTY copolymers. **a** MAASTY (left) and AASTY (right) copolymers at six different starting compositions of monomers ranging from 40-65%. X-axes represent the global monomer conversion, and y-axes represent the percentage of MAA or AA copolymerized at the given conversion^40^. All six simulated MAASTY copolymers were synthesized for this study. **b** Run lengths of MAASTY (left) and AASTY (right) at equimolar MAA or AA to STY copolymerization, with red bars showing STY run lengths, and blue bars showing MAA or AA.

**Table 1:** MAASTY copolymer characteristics. Abbreviations and symbols used: Methacrylic acid, MAA; conversion, conv.; simulated MAA fraction at given conversion, MAA frac; chemical correlation parameter, *λ* (*λ* = 0 is a perfectly random copolymer, *λ* = −1 is perfectly alternating); number-average molecular weight, M_n_; dispersity, Ð= M_w_*/*M_n_, where M_w_ is the weight average molecular weight; acid dissociation constant, pKa. ^1^H-NMR of all MAASTY copolymers is contained in Fig. S1. pKa was determined from the Bjerrum plot in Fig. S2.

| MAASTY<br>coolymers | Initial<br>MAA (%) | Conv<br>(%) | MAA<br>frac<br>(%) | $\lambda$ | $M_n$ kDa<br>$^1\text{H-NMR}$ | $M_n$ kDa<br>SEC | $\mathcal{D}$<br>SEC | pKa |
| --- | --- | --- | --- | --- | --- | --- | --- | --- |
| 7.4-40 | 40 | 81 | 44 | -0.48 | 7.4 | 5.9 | 1.68 | 8.23 |
| 7.0-45 | 45 | 73 | 48 | -0.49 | 7.0 | 3.5 | 1.55 | 7.95 |
| 7.1-50 | 50 | 75 | 51 | -0.49 | 7.1 | 8.9 | 1.99 | 7.75 |
| 8.2-55 | 55 | 88 | 55 | -0.48 | 8.2 | 5.6 | 1.69 | 7.62 |
| 8.1-60 | 60 | 88 | 59 | -0.46 | 8.1 | 6.2 | 1.35 | 7.44 |
| 8.0-65 | 65 | 88 | 63 | -0.42 | 8.0 | 6.2 | 1.74 | 7.18 |

We have previously shown that the charge state of the AASTY copolymer depends on the pH of the solution as acid groups are protonated at acidic pH values causing the polymer to become insoluble and unable to form lipid nanodiscs^18^. As MAASTY copolymers contain MAA groups, we decided to characterize the ionization state of the MAASTY copolymer library as a function of pH, identifying their respective pKa values and pH tolerances (Fig. S2; Table 1). The MAASTY copolymers displayed a single pKa in the range of 7.2-8.2. The broader polymer compositions give MAASTY copolymers an extended effective working pH range compared to AASTY copolymers with pKa values between 6.5 and 7.0 using initial AA fractions of 40%, 45%, 50% and 55%, respectively^18^. For MAASTY copolymers, the pKa decreases with increasing MAA fraction and correspondingly decreasing copolymer hydrophobicity (Table 1). This suggests that the solvation of phenyl moieties in water has a greater energetic barrier than the intra-chain charge repulsion from carboxylates. For instance, MAASTY_8.0_-65 exhibits the lowest degree of protonation at lower pH values than the other MAASTY copolymers in the library indicating that this copolymer may be more effective in solubilizing lipids and proteins at a more acidic pH compared to a higher pH. Indeed, we see that MAASTY_8.0_-65 performs better at pH 7 and 7.4 than 8 (Fig. S3). It should be noted that all experiments in this study were carried out at pH 7.4, mimicking a physiological pH; however, this may not be the optimal test conditions for the MAASTY copolymers on either end of the MAA composition spectrum.

In addition to being pH sensitive, polyanionic copolymers are chelators of divalent cations through their carboxylate moieties. This may cause complications when working with Ca^2+^ or Mg^2+^-dependent proteins in polymer-stabilized nanodiscs. Consequently, we assessed the Ca^2+^ sensitivity of MAASTY copolymers using a colorimetric detection assay of free Ca^2+^ ions with *o*-cresolphthalein complexone (oCPC), as carried out previously for AASTY^38^. oCPC yields a bright purple complex with an absorbance maximum at 575 nm upon chelating divalent cations, allowing a measurement of the free Ca^2+^ ions left in solution following copolymer exposure (Fig. S4a, b). For the tested MAASTY copolymers, the total escaped Ca^2+^ ions remained consistent at 1.2 ± 1.5 % for the 1 mM and 3 mM CaCl_2_ conditions (Fig. S4c). We measured a sudden increase to 5.1 ± 0.8 % of total escaped Ca^2+^ ions with 7 mM added CaCl_2_ for all copolymers except MAASTY_7.0_-45. This increase may occur as the copolymer becomes saturated with Ca^2+^ at this concentration. Overall, a higher absolute copolymer MAA fraction does not equate to higher Ca^2+^ binding capacity, within the concentration range tested. This is similar to what we found previously for AASTY copolymers^38^. Contrarily, the affinity of MAASTY copolymers for Ca^2+^ is seemingly higher than AASTY, given a drastically different 21% of total escaped Ca^2+^ ions was measured for equivalent assayed concentrations in the presence of AASTY^38^. This lower Ca^2+^ tolerance of MAASTY compared to AASTY may be due to its increased hydrophobicity, resulting in Ca^2+^ being kinetically trapped within solubilized aggregates of MAASTY-Ca^2+^ complex.

To further understand the impact of divalent cations on the nanodisc-forming ability of MAASTY copolymers, we produced fluorescent nanodiscs by solubilizing small unilamellar vesicles (SUVs) composed of 1-palmitoyl-2-oleoyl-*sn*-glycero-3-phosphocholine (POPC) and 2% fluorescent 1,2-dioleoyl-*sn*-glycero-3-phosphoethanolamine-N-(lissamine rhodamine B sulfonyl) (LissRhod) phosphatidylethanolamine (PE) with MAASTY copolymers in the presence of specific CaCl_2_ and MgCl_2_ concentrations and subjected them to FSEC (Fig. S5). As previously exhibited for SMA and AASTY copolymers^27,38^, solubilization by MAASTY copolymers is more sensitive to Ca^2+^ ions than Mg^2+^ ions. Additionally, as divalent cation concentrations increase, the fraction of larger soluble species also increases, a phenomenon that may be caused by rouleaux stacking of nanodiscs under these conditions^42^. In contrast to the free copolymer assessed in the oCPC assay (Fig. S4), the tolerance towards Ca^2+^ and Mg^2+^ ions during nanodisc-formation appears to depend on copolymer composition. Of note, MAASTY copolymers with higher MAA content than STY (MAASTY_8.2_-55, MAASTY_8.1_-60 and MAASTY_8.0_-65) were observed to better tolerate the presence of divalent cations during nanodisc-formation, solubilizing lipids either as well or to a greater extent in the presence of Ca^2+^ or Mg^2+^ ions (Fig. S5g).

### MAASTY facilitates high-resolution structural determination of integral membrane proteins

To benchmark the utility of MAASTY copolymers for structural determination of integral membrane proteins, we screened the solubilization capacity of our MAASTY copolymer library on full-length hTRPM4, N-terminally tagged with StrepTag-IIeGFP, and recombinantly expressed in HEK293 cells with FSEC (Fig. S6) and compared it to detergent-solubilization (Fig. S7). hTRPM4 is a Ca^2+^-activated, non-selective monovalent cation channel that is broadly expressed and contributes to the pathophysiology of cardiovascular diseases, prostate cancer, and diseases of the central nervous system^43^. Notably, solubilization with MAASTY outperforms DDM-CHS (Fig. S7). In accordance with FSEC screening of the MAASTY copolymer library, hTRPM4 was solubilized and purified in MAASTY_7.1_-50 nanodiscs (Fig. S6; S8; S9), and subjected to single-particle cryo-EM analysis, resulting in a structure with an overall resolution of 3.5 Å (Fig. 3a; Fig. S10; S11; Table S1). The transmembrane domain and extracellular loops, the N-terminal, cytosolic TRPM homology region 4 (MHR4) and the central coiled-coil in the C-terminus were well-resolved, and overall identical to previously determined cryo-EM hTRPM4 structures in detergent micelles or MSP nanodiscs^44–47^. Despite extensive 3D classifications and focused refinements imposing either C1 or C4 symmetry, the portions of the cytosolic domain composed by the MHR1-3 of hTRPM4 in MAASTY nanodiscs were unresolved, likely due to conformational flexibility under the experimental conditions (Fig. 3b). Structural flexibility of this part of the cytosolic domain is also evident from the structure of hTRPM4 in MSP2N2 nanodiscs performed in parallel with hTRPM4 in MAASTY nanodiscs (Fig. S12; S13) and as previously described^45^, albeit not as severe as in MAASTYstabilized nanodiscs. Notably, a recent structure of hTRPM4 in SMA nanodiscs also resulted in an unresolved cytosolic region of MHR1-2^21^, underscoring that hTRPM4 is more flexible in copolymer-based lipid nanodiscs, possibly due to larger heterogeneity in the lipid composition compared to the reconstituted MSP nanodisc system and detergent micelles or, in part, due to differences in lipid packing protection provided by the nanodisc-stabilizing copolymers or proteins.

**Fig. 3:**
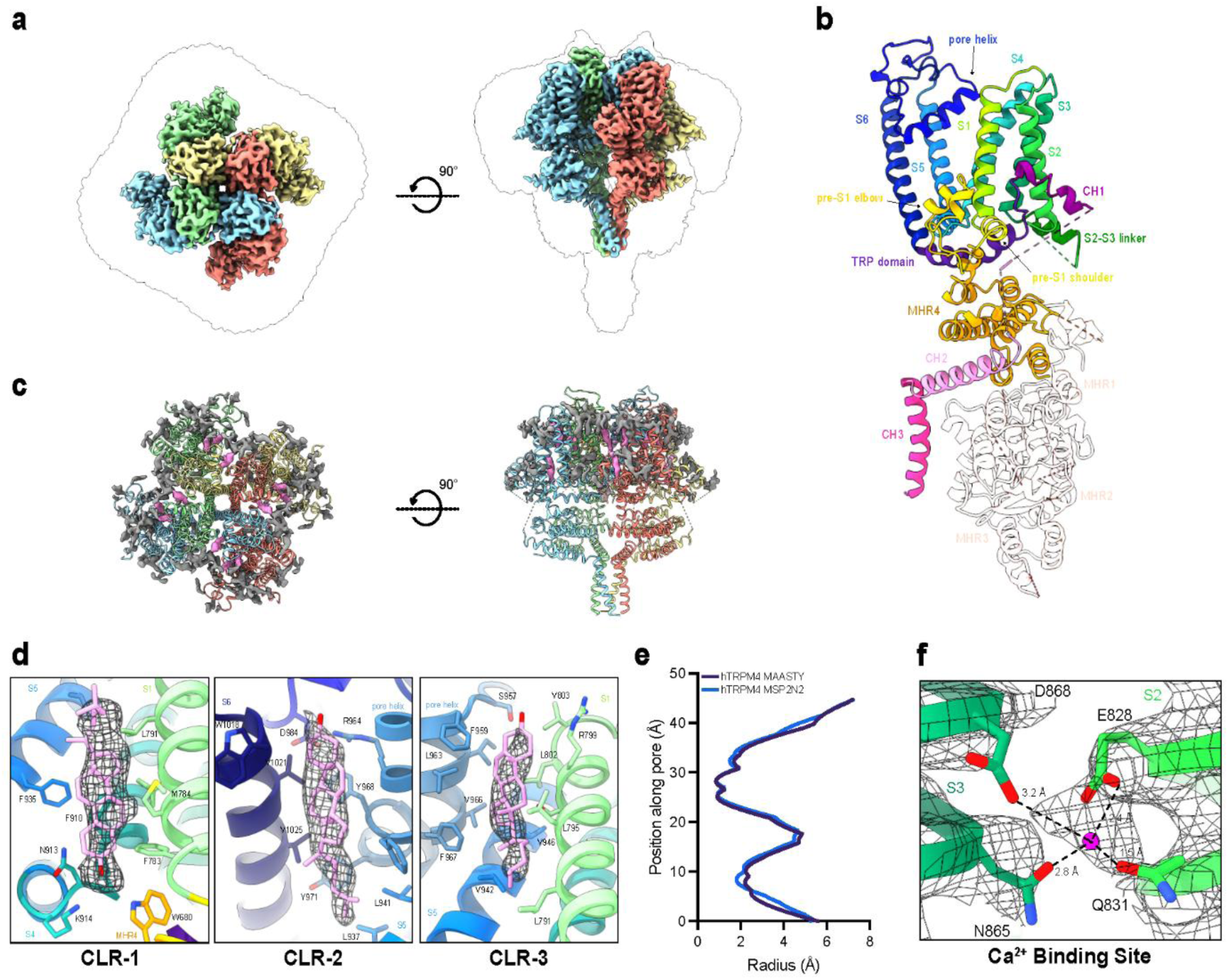
Structure of hTRPM4 in MAASTY native lipid nanodiscs. **a** Top and side views of sharpened cryo-EM density maps of hTRPM4 in MAASTY nanodiscs with the four protein subunits colored in light green, salmon, sky blue and khaki overlaid with the unsharpened cryo-EM density map (black outline). The unsharpened map represents the raw reconstruction, showing the full range of densities, including weaker and flexible regions such as copolymers and lipids in the nanodiscs. The sharpened map has been processed with a B-factor of −92 Å² to enhance high-resolution features, improving clarity in well-ordered regions. Contour levels for both maps were adjusted for optimal visualization of key structural features. **b** Atomic model of the monomeric hTRPM4 in MAASTY nanodiscs in ribbons with domains labelled and separately colored. Unresolved domains of MHR1-3 are displayed as transparent ribbons using the atomic model of monomeric hTRPM4 in MSP2N2 nanodiscs. **c** Top and side views of resolved lipid densities among the transmembrane domain of hTRPM4 in MAASTY. The atomic model of hTRPM4 is represented as ribbons and colored as in **a**. Lipid densities are shown in grey and densities characterized as CLR molecules are shown in pink. **d** Close-up views of the positions of identified CLR molecules: CLR-1, CLR-2 and CLR-3, identified in hTRPM4-MAASTY structure. The cryo-EM map densities are represented as mesh and modelled CLR molecules shown as licorice in pink. Domains are labelled and colored as in **c**. **e** Pore profiles of hTRPM4-MAASTY (purple) and hTRPM4-MSP2N2 (blue) structures exhibiting similarly closed states. **f** Identified Ca^2+^ binding site shown as coordination of the bound Ca^2+^ ion by residues Glu^828^, Gln^831^, Asn^865^, and Asp^868^. Distances are labelled and domains denoted as in **c**.

From analyzing the hTRPM4-MAASTY structure, we found several extra densities that we identified as lipids (Fig. 3c). Along with observing densities of lipids, we modelled three densities as cholesterol (CLR) molecules due to its distinct shape and location (Fig. 3d). The identified CLR molecules in site 1 and site 2 (CLR-1 and CLR-2, respectively) lie within sites previously annotated as cholesteryl hemisuccinate (CHS) binding sites by Autzen *et al.*^45^, and discerned in the hTRPM4-MSP2N2 structure of this study (Fig. S12; S13). CLR-1 is located at the pre-S1 elbow, creating a cavity with S1, S4 and MHR4 helix 7, and CLR-2 lines the pore, interacting with the pore helix and S6, supposedly stabilizing the pore conformation^45^. CLR site 3 (CLR-3) is situated between S1, S5 and the pore helix of the adjacent proteomer, and was also observed in the structure of hTRPM4 in SMA nanodiscs^21^. Both hTRPM4 structures determined in SMA^21^ and MSP2N2^45^, including this study, identified CLR or CHS in a binding site that corresponds to the vanilloid binding pocket of TRPV1^48^. We also observed a similar density at that location in the hTRPM4-MAASTY structure, however due to low occupancy, we were unable to definitively model it as CLR.

In order to assess the endogenous lipid species co-purified with hTRPM4 in MAASTY native nanodiscs, we extracted the lipids from the purified sample and analyzed it with MALDI-TOF mass spectrometry (MS). This confirmed the presence of CLR in the nanodiscs, in addition to POPC (Fig. S14). As a further, in-depth evaluation of the relative abundance of the extracted lipid species, we subjected the hTRPM4-MAASTY nanodisc sample to lipidomics analysis. This identified a variety of putative endogenous lipid species co-extracted with hTRPM4 (Fig. S15), including lipids belonging to the phospholipid classes phosphatidylinositol (PI), phosphatidylcholine (PC), phosphatidylethanolamine (PE), phosphatidylglycerol (PG) and phosphatidylserine (PS). PE, PI and PC constituted a majority of the identified lipid classes. The lipidomics analysis also detected plasmenyl-, plasmanyl- and lysophospholipids, highlighting the complexity of membrane lipid species stabilizing, and possibly regulating, hTRPM4 in an endogenous lipid environment.

Interestingly, the MAASTY nanodisc encapsulating hTRPM4 exhibits a distinct shape. When looking down the ion channel pore axis, the nanodisc is observed as a ”square” with an overall shape similar to the transmembrane domain of hTRPM4 itself, rather than circular as for hTRPM4 in MSP2N2 nanodiscs (Fig. 3a; S12b)^45^. This resolved nanodisc shape was also observed for hTRPM4 in SMA nanodiscs^21^, appearing as an extension of the shape formed by the TM domains themselves, and likely derived from the higher degrees of freedom in flexibility of the copolymer compared to, for example, the rigidity of MSP helices. This is an attribute shared by other resolved cryo-EM structures of copolymer-encapsulated nanodiscs^16,49,50^. Despite the discrepancy in nanodisc shapes, the transmembrane domains were identical between the hTRPM4 structures in MAASTY and MSP2N2 nanodiscs with matching pore profiles (Fig. 3e).

Next, we sought to estimate the composition of the hTRPM4-MAASTY particles. The mass contribution from the individual components in the nanodisc is apparent when assessing the sample by mass photometry (MP) and spectroscopic quantification of hTRPM4 and MAASTY copolymers in the nanodiscs. MP of hTRPM4 in MAASTY revealed a peak around 760 kDa (Fig. S8c). Subtracting the mass of hTRPM4 (540 kDa for the cleaved tetramer), the excess mass of approximately 220 kDa may be attributed to extracted lipids and surrounding copolymers of the nanodisc. To assess the composition of the solubilized particles, we solubilized and purified hTRPM4 in MAASTY_7.1_-50 with a trithiocarbonate (ttc) moeity at the terminal, which has a maximum absorbance at 310 nm, and quantified hTRPM4-MAASTY_7.1_-50_ttc_ with FSEC and the Bradford protein assay (Fig. S8d, e). We estimate that for each solubilized particle, the copolymer contributes approximately 110 kDa, with 110 kDa remaining as contribution from lipids. At the molecular level, this means that every hTRPM4-MAASTY particle contains an average of 13 MAASTY copolymers.

Finally, from analysis of the hTRPM4-MAASTY structure, we observed an apparent density representing a Ca^2+^ ion in the established Ca^2+^-binding site (Fig. 3f), coordinated by side chain groups of Glu^828^, Gln^831^, Asn^865^ and Asp^868^ located at the S2-S3 helices^44,45,47^. This observation indicates that hTRPM4 in MAASTY nanodiscs can capture endogenous Ca^2+^ without the need to supplement it throughout purification or cryo-EM sample preparation. Moreover, it lends promise to the potential use of MAASTY copolymers to characterize divalent cation-bound membrane proteins despite being polyanionic.

### MAASTY solubilizes a diverse test set of membrane proteins

Next, we assessed the the versatility of the MAASTY copolymers across varying protein types through the solubilization of three additional eukaryotic integral membrane proteins, fused with eGFP, and recombinantly expressed in HEK293 cells. The test set included three differing oligomeric configurations, represented by the rabbit sarco/endoplasmic reticulum Ca^2+^-ATPase 1a (rSERCA1a, monomeric), the human potassium channel subfamily K member 18 (hKCNK18, homodimeric), and the chicken acid sensing ion channel 1 (cASIC1, homotrimeric), in addition to the homotetrameric hTRPM4 presented in the previous section (Fig. S6). For each membrane protein, we compared the solubilization efficacy of the MAASTY copolymer library with a selection of six AASTY copolymers with initial AA content of 45%, 50% and 55% and masses of either 6 kDa or 11 kDa, respectively (Fig 4; Table S2; Fig. S16). rSERCA is a P-type ATPase that pumps Ca^2+^ ions from the cytoplasm into the sarcoplasmic or endoplasmic reticulum, and plays a critical role in muscle contraction and cellular Ca^2+^ homeostasis^51^. With 79 deposited models in the PDB, rSERCA is structurally and functionally well-characterized.

**Fig. 4:**
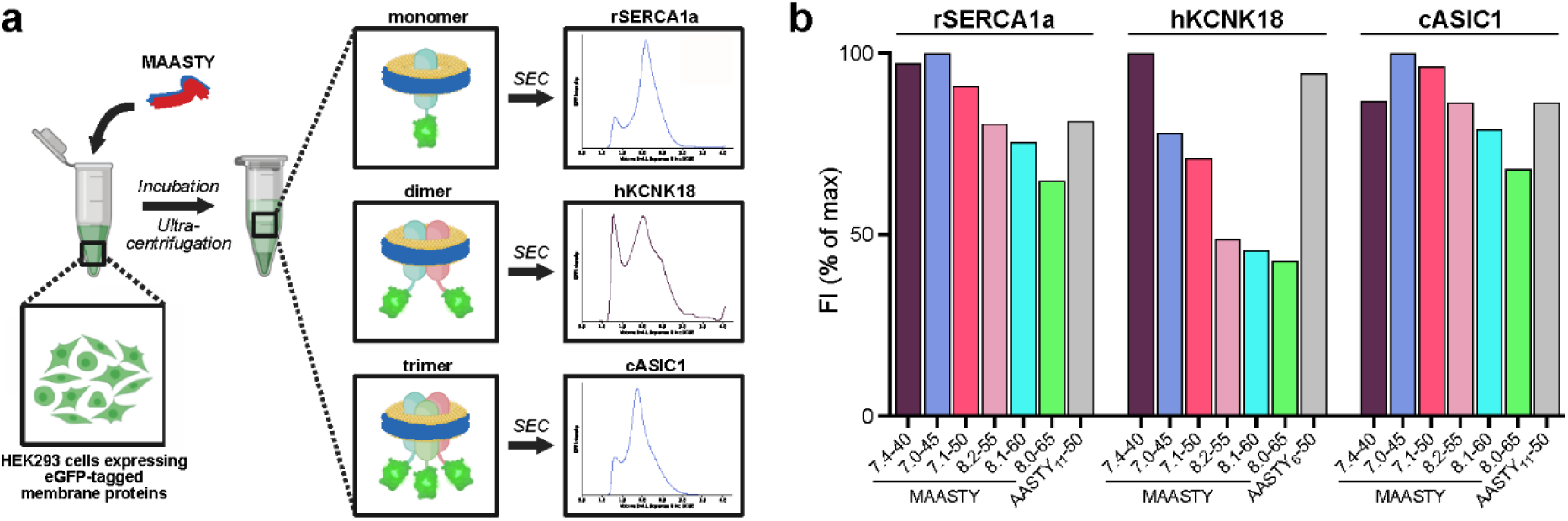
Solubilization of eukaryotic membrane proteins by MAASTY copolymers. **a** Schematic of preparation of MAASTY-solubilized integral membrane proteins from HEK293 cell lysate. First, 2% MAASTY copolymer is added to HEK293 cells recombinantly expressing eGFP-tagged membrane protein and incubated to enable formation of protein-loaded nanodiscs. Insoluble material is then removed via ultra-centrifugation and the soluble fraction is assessed by FSEC using an excitation wavelength of 488 nm and emission of 507 nm. In this study, we assessed the solubilization efficiency of MAASTY copolymers using a test set of membrane proteins including monomeric rSERCA1a, homodimeric hKCNK18, and homotrimeric cASIC1. The displayed example FSEC traces for each test protein (box) represent those with the highest peak intensity at the presumed proteinnanodisc peak. All FSEC traces from all tested copolymers for each membrane protein is found in Fig. S16. Each unique membrane protein copolymer sample was injected twice or more to check reproducibility. **b** Normalized maximum fluorescence intensity values of the protein peaks from the FSEC traces of each tested copolymer to compare differences in solubilization efficiencies. For each protein, the performance of MAASTY copolymers are compared to the best performing AASTY copolymer, as determined by the maximum fluorescence intensity of the presumed protein-nanodisc peak. The maximum fluorescent intensity values of all tested copolymers for each membrane protein is found in Fig. S16.

Solubilization of rSERCA1a with both MAASTY and AASTY copolymers, produced a largely monodisperse peak with a trailing shoulder at longer retention times (Fig. S16a). MAASTY_7.4_-40 and MAASTY_7.0_-45 were most effective at solubilizing rSERCA1a, with decreasing solubilization efficiency as the MAA content of the copolymer increased, with two of six MAASTY copolymers performing worse than the best AASTY copolymer (Fig. 4b; Fig. S16a). While most of the copolymers showed the same retention times, three of the MAASTY copolymers, MAASTY_8.2_-55, MAASTY_8.1_-60, and MAASTY_8.0_-65 showed slightly earlier retention times, which may be indicative of larger discs compared to the later eluting species (Fig. S16a).

hKCNK18 is a potassium channel implicated in regulation of neuronal excitability and associated with certain types of migraines^52^. The structure of hKCNK18 is yet to be determined, probably due to difficulties in its expression and stabilization outside the cell membrane. For hKCNK18, all MAASTY copolymers led to FSEC traces with shoulders, making it difficult to estimate the solubilization efficacy and the retention volumes (Fig. S16b). MAASTY_7.0_-40 seemingly displayed the highest solubilization capacity, producing a more uniform peak than the five other MAASTY copolymers and any of the tested AASTY copolymers (Fig 4b; Fig. S16b).

cASIC1 is a proton-gated Na^+^ channel involved in detection of acidic environments. With approximately 90% sequence identity to human ASIC1a, a protein implicated in a variety of neuropathologies^53^, cASIC1 is the most structurally well-characterized of the ASICs amounting to 19 deposited models in the PDB. A majority of these were determined in detergent micelles, but some were resolved in SMA^19^. For cASIC1, both MAASTY and AASTY copolymers acted as effective solubilizers, generating a uniform presumed protein-nanodisc peak with a trailing shoulder on the right side of the main peak (Fig. S16c).

MAASTY_7.0_-45 and MAASTY_7.1_-50 were the most potent solubilizers, however, most of the MAASTY copolymers solubilized cASIC almost as well (Fig. 4b). As with rSERCA1a, four of six MAASTY copolymers produce higher peak intensities than the best-performing AASTY copolymer. Interestingly, there was a distinct difference in retention volumes between MAASTY and AASTY copolymer nanodiscs for cASIC1, with MAASTY copolymers eluting at earlier volumes forming evidently larger cASIC1-containing nanodiscs than AASTY copolymers (Fig. S16c). This may also be the case for hKCNK18, indicating a possibility that MAASTY copolymers form larger protein-containing nanodiscs than AASTY copolymers.

### MAASTY is compatible with a broad selection of lipids

As previously described, the efficiency of nanodisc formation by AASTY copolymers is highly dependent on membrane charge, specifically favoring more zwitterionic lipid compositions^18^, whereas the anionic SMA and DIBMA copolymers reportedly solubilize irrespective of lipid charge^54–56^. To assess the effect of lipid composition on MAASTY solubilization, we dissolved 2% fluorescent LissRhod PE and one of three lipid mixtures: POPC, soy polar lipid extract (PLEx) and *E. coli* PLEx with MAASTY copolymers to form protein-free or ”empty” nanodiscs. Using FSEC, we determined the lipid solubilization capacity for each MAASTY copolymer from the fluorescence intensity (Fig. 5; Fig. S17). All six MAASTY copolymers formed nanodiscs with all three lipid compositions, and although the efficiencies vary, the MAASTY library forms nanodiscs with both the synthetic lipid POPC and the two lipid extracts (Fig. 5b). This is in contrast to AASTY, which showed significantly decreased efficiency for soy PLEx and *E. coli* PLEx, likely owing to their content of charged lipid species^18^. From the MAASTY library, MAASTY_7.0_-45 and MAASTY_7.1_-50 were the most effective copolymers for all three of the tested lipid compositions. In contrast, MAASTY_8.0_-65, which is the copolymer with the highest MAA fraction (63%), was the least effective at solubilizing the three lipid compositions. MAASTY_8.0_-65 was particularly poor at solubilizing *E. coli* PLEx, which has a high content of negatively charged lipid species such as cardiolipin (∼10%) and phosphatidylglycerol (∼23%); This could be a charge effect as MAASTY_8.1_-60, with the second highest MAA fraction (59%), also displayed a diminished capacity to solubilize *E. coli* PLEx (Fig. 5b). Indeed, with pKa values at 7.2 and 7.4, respectively, these two copolymers are likely either predominantly charged or at least 50% charged at pH 7.4 (Table 1; Fig. S2).

**Fig. 5:**
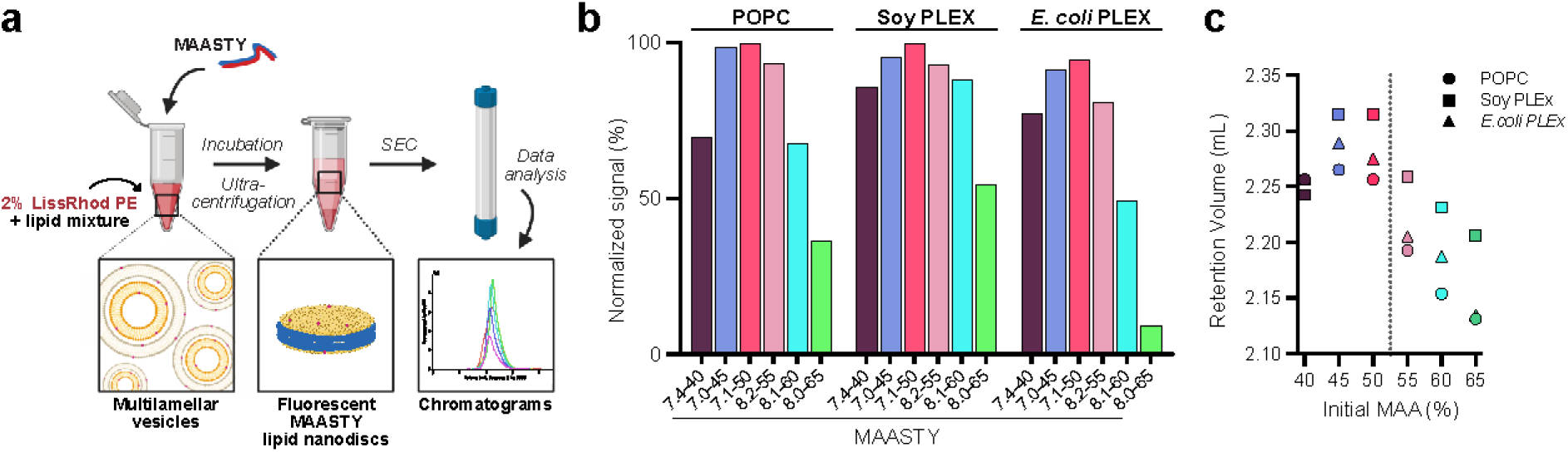
Solubilization of lipids into nanodiscs by MAASTY copolymers. **a** Schematic of preparation of empty native nanodiscs from hydrated lipid films, illustrating the process. First, 2% LissRhod PE is added to a defined lipid composition forming multilamellar vesicles. MAASTY copolymer is added and the mixture and incubated to allow for solubilization. Next, insoluble material is removed via ultra-centrifugation, while the remaining soluble material is assessed by FSEC using an excitation wavelength of 554 nm and emission of 576 nm. Finally, resulting FSEC traces of the tested MAASTY copolymers for differing lipid compositions are compared. **b** Max peak values of the MAASTY-forming empty nanodisc FSEC traces with POPC, Soy PLEx and *E. coli* PLEx mixed with 2% Liss Rhod PE from Fig. S17. Values are normalized relative to the highest detected signal for the contained data. **c** The retention volumes determined from the max peak values of the MAASTY-forming empty nanodisc FSEC traces with POPC, soy PLEx and *E. coli* PLEx from Fig. S17 compared to the initial MAA content (%) of the tested MAASTY copolymers.

Notably, we observed a disparity in retention volumes across the tested MAASTY copolymers and lipid compositions (Fig. 5c). Overall, there appears to be a trend of lower retention volumes with increasing copolymer MAA content. For example, MAASTY_8.0_-65, which has the highest MAA fraction of all tested copolymers, has the smallest retention volumes, particularly when forming nanodiscs of POPC and *E. coli* PLEx. In general, POPC and *E. coli* PLEx containing nanodiscs display lower retention volumes than soy PLEx for each of the MAASTY copolymers, except MAASTY_7.4_-40. Whether the dissimilarity in retention volume is an indication of differences in the relative disc sizes, i.e. a creation of larger nanodiscs from lower retention volumes, or rather a result of the charge state of the polymer, remains to be assessed further.

## Discussion

The limited number of structural studies of membrane proteins in native nanodiscs underscores a requirement to improve our current repertoire of nanodisc-forming copolymers. Only recently did the first high resolution structures of human membrane proteins in native nanodiscs emerge, emphasizing the potential of the method^21,22^. Here we introduce the copolymer MAASTY, which can solubilize membrane proteins in native nanodiscs and facilitate high-resolution structural determination. We previously conceptualized AASTY as an effective copolymer for formation of native nanodiscs, based on the structural regularity of this copolymer^18^. Since then, it was found that altering the hydrophobicity of the styrenic moiety of alternating SMA copolymers also yields effective nanodisc forming copolymers^57^. On this basis, we envisioned that changing AA to MAA could result in an improvement, as the methyl group of MAA is more hydrophobic. Previous research into nanodisc forming copolymers shows that the main properties determining nanodisc-forming capabilities are the amphiphilicity, size, monomer sequence and monomer composition of the copolymers^57–60^. Delineating the effects of these four properties on nanodisc formation is non-trivial as exchanging a monomer with another results in a different statistical distribution of monomers along the chain (Fig. 2). Our rationale for designing AASTY and our hypothesis on why it is an effective solubilizer was that the alternating monomer sequence, in combination with a similar amphiphilicity to effective SMA compositions, would be more effective than SMA due to the high regularity in monomer sequence^18^. As such, we were somewhat surprised that MAASTY, which is more irregular than AASTY, proved to be as effective in nanodisc-formation and, for some targets, seemingly performed better than AASTY. However, by using MAA instead of AA, we change both the amphiphilicity and the statistical distribution of monomers in the polymer, emphasizing that observed differences between MAASTY and AASTY, or other copolymers for that matter, with regard to nanodisc formation cannot be attributed to one particular polymer property. The key difference of MAASTY to AASTY is the increased hydrophobicity of MAASTY, and it could be that the packing of the methyl group into the lipid bilayer is beneficial for the stabilization of nanodiscs. As polymers with methylated backbones have lower glass transition temperatures than their non-methylated acrylate counterparts, this feature could result in a more rigid structure, conferring higher stability of the disc. The more hydrophobic nature of the MAASTY copolymers will also likely result in increased sensitivity to divalent cations, at least for the free copolymers, as the charge of the carboxylates are neutralized by Ca^2+^ ions, resulting in precipitation. We hypothesize that this is the reason for the pronounced Ca^2+^ ion binding we see with MAASTY copolymers in the oCPC assay (Fig. S4). Furthermore, the large span of pKa values for the MAASTY copolymer library synthesized in this study is a result of the wider range of compositions enabled by the less-alternating nature of the copolymerization in comparison to AASTY (Table 1). More MAA in a copolymer results in a markedly lower pKa, which can only mean that the energy penalty for deprotonating MAA afforded by the local hydrophobic environment of STY moieties is larger than the penalty from adjacent negative charges from deprotonated carboxylates. In practice however, this likely means that MAASTY copolymers can be tailored to perform optimally at a desired pH.

The high-resolution cryo-EM structure of hTRPM4 in MAASTY native nanodiscs showcases the applicability of MAASTY copolymers and expands the toolbox for membrane protein characterization (Fig. 3; 4). The structure elaborates on our previous research on AASTY copolymers in which sample and conformational heterogeneity yielded low-resolution reconstructions of hTRPM4 in native nanodiscs^18^. There may be many possible explanations for the successful use of MAASTY for high-resolution structural determination of hTRPM4 by cryo-EM in comparison to AASTY, such as altered orientational distribution, number of particles at the air-water interface and conformational flexibility resulting from differences in overall charge of the protein-nanodisc complex. In addition, we cannot exclude that a high resolution structure with AASTY may have been feasible to obtain from a larger dataset. It is also possible that potential differences in the co-purified lipids between AASTY and MAASTY contribute to the success of MAASTY for the structural determination of hTRPM4. Indeed, lipid solubilization with MAASTY and AASTY show different lipid preferences for the two copolymers; AASTY displayed a lower propensity to solubilize negatively-charged lipids^18^, while MAASTY solubilizes lipids independently of head-group charge (Fig. 5).

According to lipidomics analysis of hTRPM4-MAASTY nanodiscs, two of the top five most abundant lipid species co-extracted with hTRPM4 are the negatively charged lipid species PI and PS (Fig. S15). Notably, with an observed relative abundance of 15-25%, PI co-purified in hTRPM4-MAASTY nanodiscs far exceeds the typical plasma membrane content of 1-2%^61^. Whether the enrichment of PI and PS is due to hTRPM4, the HEK293 cells or the MAASTY copolymers should be further tested. Along with its reported role in channel regulation, the phosphorylated PI, phosphatidylinositol 4,5-bisphosphate (PIP2), has been hypothesized to aid in conformationally stabilizing hTRPM4^62,63^. We speculate that differences in extraction of charged lipids between AASTY and MAASTY copolymers may have resulted in less co-purified PI in the hTRPM4-AASTY sample, along with other negatively-charged lipids, consequently displaying increased conformational heterogeneity and potentially contributing to the yielding of a lower resolution structure, among other aformentioned possibilities. Furthermore, other lipids such as CLR may be necessary to stabilize hTRPM4^18,44–46,64^. The stability of the transmembrane domain by lipids may also play a role in the missing cytosolic MHR1-3 domains in the hTRPM4-MAASTY structure which we assume is caused by structural flexibility. This structural flexibility could even be potentiated by the copolymers themselves either through direct or indirect interactions with the protein. Indeed, this has been reported for membrane proteins in MSP nanodiscs whereby amphipathic interactions with the MSP may influence the local dynamics of the membrane protein^65^. Lastly, MAASTY copolymers consistently solubilize hTRPM4 to a greater extent than AASTY copolymers, with MAASTY_7.1_-50 as the most efficacious, potentially contributing to better sample quality (Fig. S6). hTRPM4 solubilization by MAASTY_7.1_-50 was even shown to outperform DDM-CHS (Fig. S7).

Endogenous lipids are clearly visible in the hTRPM4-MAASTY structure and further confirmed by lipidomics analyses (Fig. 3; S14; S15). The observed arrangement of lipid densities in the hTRPM4 MAASTY structure differed from that of hTRPM4 in MSP2N2. Specifically, we identified the locations of three CLR binding sites; two of which are also occupied in the hTRPM4-SMA structure^21^, and a novel location previously annotated as a CHS-binding region^45^, adding to our current knowledge on lipid organization in hTRPM4. MS analyses highlighted the complexity of the co-purified endogenous lipid species unassigned in the hTRPM4-MAASTY structure. These identified lipids may be implicated in hTRPM4 activity, regulation, and/or stabilization. For example, the detection of lysophospholipid classes could indicate a potential role in channel modulation, as established in other TRP channels^66,67^.

In addition to endogenous lipids, hTRPM4 in MAASTY nanodiscs were bound to endogenous Ca^2+^ ions, according to an observed density in the Ca^2+^-binding site without having supplemented the sample with CaCl_2_ during purification or cryoEM sample preparation. The binding of Ca^2+^ from an endogenous source was also eluded to in the hTPRM4-SMA structure^21^, while this was not reported for any of the detergent-solubilized structures^44–46,64^. The ability to capture a Ca^2+^-bound state of hTRPM4 gives evidence towards the use of MAASTY copolymers to characterize divalent cation sensitive membrane proteins despite the polyanionic nature of MAASTY and consequent propensity to bind Ca^2+^. Indeed, we show that MAASTY-encasing protein-free nanodiscs are tolerant of CaCl_2_ and MgCl_2_ (Fig. S5). These observations underscore the importance of assessing tolerance to solvent compositions, such as the addition of divalent cations, under nanodisc-forming conditions, as opposed to solely measuring the impact on free copolymer (Fig. S4). While the enhanced tolerance of higher MAA content copolymers suggest they may be particularly well suited for studying Ca^2+^ or Mg^2+^ sensitive proteins, this is likely also influenced by the properties of the solubilized membrane protein. In general, the decrease in accessible divalent cations by copolymer chelation, and consequent concentration change throughout membrane protein purification, are both essential effects to be aware of.

Finally, we demonstrate that MAASTY is a versatile copolymer capable of solubilizing a range of disease-relevant membrane proteins and diverse lipid compositions (Fig. 4; 5). Interestingly, we noted differences in SEC elution volumes between MAASTY and AASTY nanodiscs incorporating membrane proteins, as well as retention volume differences for MAASTY-encasing protein-free nanodiscs of increasing negatively-charged lipid compositions. These observations indicate that solubilization efficacy is dependent on the membrane protein solubilized, the lipid species, and the copolymer itself.

While it is unlikely that there is a one-size-fits-all solution of nanodisc-forming copolymers, MAASTY is an easily synthesized copolymer, able to yield a high resolution structure in a nanodisc without the need for extensive reconstitution optimization. Understanding the mechanism of how MAASTY and other copolymers work will ease the identification of which copolymers are effective for specific protein targets in a given membrane composition. Further lipidomics analyses of native nanodiscs could help elucidate how lipids influence other target-copolymer combinations. We conclude that our work showcases the potential of MAASTY in membrane protein biochemistry and structural biology, and lends promise to its further use in the field as an effective solubilizer of integral membrane proteins in native lipid nanodiscs across varying recombinant-expression and native systems.

## Methods

### Synthesis and characterization of MAASTY copolymers

Methacrylic acid (MAA) and styrene (STY) were distilled under vacuum. The RAFT agent 2-Methyl-2-[(dodecylsulfanylthiocarbonyl)sulfanyl]propanoic acid (194 mg, 0.531 mmol) and azobisisobutyrunitrile (17.4 mg, 0.106 mmol) were added to six ampules. Styrene and methacrylic acid were added to the ampules; MAASTY_7.4_-40 (MAA: 1.7 mL, 1.7 g, 20 mmol, STY: 3.44 mL, 3.13 g, 30 mmol), MAASTY_7.0_-45 (MAA: 1.91 mL, 1.94 g, 22.5 mmol, STY: 3.15 mL, 2.85 g, 27.5 mmol), MAASTY_7.1_-50 (MAA: 2.12 mL, 2.15 g, 25 mmol, STY: 2.86 mL, 2.60 g, 25 mmol), MAASTY_8.2_-55 (MAA: 2.33 mL, 2.37 g, 27.5 mmol, STY: 2.58 mL, 2.34 g, 27.5 mmol), MAASTY_8.1_-60 (MAA: 2.55 mL, 2.60 g, 30 mmol, STY: 2.29 mL, 2.10 g, 20 mmol), MAASTY_8.0_-65 (MAA: 2.76 mL, 2.80 g, 32.5 mmol, STY: 2.05 mL, 1.82 g, 18.5 mmol). The ampules were degassed by four freeze-pump-thaw cycles, and sealed under vacuum while frozen. They were then placed in a water bath in a 60 °C oven for 17h, forming yellow solids. The solids were removed by ampule shattering and each dissolved in 10 mL absolute ethanol. At this point, samples for determining conversion by NMR were taken. 20 equiv. of H_2_O_2_ (30% in water) was added with respect to end-groups. These mixtures were heated to 70 °C overnight resulting in non-colored solutions. The solutions were precipitated into heptane and the white precipitate was recovered by filtration. The polymers were converted to their respective sodium salts by suspending them in ultrapure water and titrating in 1 M NaOH until the pH was stable at approximately pH 8.0, and the bulk of the suspended polymer was dissolved. These solutions were filtered through 0.22 µM PES syringe filters and freeze-dried.

Bjerrum diagrams were constructed by dissolving 50 mg of respective polymer in 10 mL water and adjusting the pH to approximately 11 with 1 M NaOH. 0.33 M HCl solution was titrated in 5 µL increments whilst measuring the pH on a Mettler Toledo SevenCompact S210 pH meter.

We were unable to use aqueous SEC Multi-Angle Light static Scattering (MALS) to characterize the MAASTY copolymers. Even at minimal concentration, MALS would report masses that were orders of magnitude higher, than expected for the given elution time. Expecting these effects were due to micellization, we performed the analysis in organic SEC, after methylation, as previously described^68^. The analysis showed a greater dispersity than what could be expected for RAFT made polymers. We believe that this is due to incomplete methylation resulting in aggregation or crosslinking of polymers.

### NMR characterization of MAASTY

Conversion of monomers were determined by ^1^H-NMR using a Bruker Avance III 400 MHz system using d6-DMSO and a probe temperature of 25 °C. The ^1^H-NMR of the purified polymers converted to sodium salts were obtained in D_2_O (Fig.S1).

### Methylation and SEC characterization of MAASTY polymers

MAASTY polymer (1 equiv.) was suspended in a 1 w% 1-methyl-3-*p*-tolyltriazene (MTT) solution (1.2 equiv.) in toluene. The mixture was left to stir overnight. The reaction was finished when all polymer had dissolved in the toluene. Once the reaction was finished, the toluene was washed twice with 1 M HCl and the solvent was removed under vacuum. The degree of methylation was confirmed with NMR. Methylated MAASTY was dissolved in THF to a concentration of 2.5 mg/mL. Measurements were performed on a Viscotek Differential Refractometer viscometer with SIL-10AD VP Shimadzu Autosampler and LC-10ADVP Shimadzu Pump using THF as solvent and PLgel 3um MIXED E Column of 300 mm x 7.5 mm cross-linked porous polystyrene divinylbenzene matrix from Polymer Laboratories.

### Determination of free Ca^2+^ concentrations

To measure the concentration of free Ca^2+^ ions in the presence of MAASTY, copolymers were mixed with buffer for a final composition of 20 mM HEPES/NaOH (pH 7.4), 100 mM NaCl, and 0, 1, 3, or 7 mM CaCl_2_ and either 0.1% or 1% weight per volume (w/v) copolymer. After 15 min of incubation, ∼10 µL of buffer without copolymer was isolated using a Vivaspin-500 centrifugal concentrator with a 10 kDa MWCO cutoff. In a 96-well plate, 5 µL of this solution was then mixed with 95 µL of 0.13 mg/mL *o*-cresolphthalein complexone (oCPC) in 0.1 M CAPS, pH 10. The absorbance at 575 nm was measured on a Tecan Infinite M200 pro microplate reader and quantified on the basis of a standard CaCl_2_ curve (Fig. S4c).

### Preparation of protein-free lipid nanodiscs

Preparartion of protein-free lipid nanodiscs was carried out as described by Smith *et al.*^18^ and Timcenko *et al.*^38^. In short, dried lipids were hydrated in HBS (50 mM HEPES/NaOH (pH 7.4 unless otherwise stated) and 150 mM NaCl) to 10 mM final concentration for the tested lipid mixtures: 1-palmitoyl-2-oleoyl-*sn*-glycero-3-phosphocholine (POPC); soy polar lipid extract (Soy PLE); *E. coli* polar lipid extract (*E. coli* PLEx), which were mixed with 0.12 mM fluorescent 1,2-dioleoyl-*sn*-glycero-3-phosphoethanolamine-N-(lissamine rhodamine B sulfonyl) (LissRhod PE). All lipids were purchased from Avanti Polar Lipids Plc. Empty fluorescent nanodiscs were then prepared by mixing 1% MAASTY copolymer with 1 mM total lipids (small unilameller vesicles (SUVs) composed of 2% LissRhod PE and 98% chosen lipid mixture) in HBS with or without specified CaCl_2_ or MgCl_2_ concentrations in a 50 µL total volume. The mixture was incubated on rotation for 2 h at 4 °C, followed by ultracentrifugation at 113,700 xg for 15 min at 4 °C to remove insoluble material. The lipid nanodisc mixture was filtered through a 0.22 µm spin-filter before injection onto a Superose 6 Increase 5/150 GL column (Cytiva) equilibrated in HBS at 4 °C. The eluent was detected by a Shimadzu (Shimadzu Europa GmbH) liquid chromatography system equipped with an autosampler (SIL-40), a fluorometer (RF-20A), and a PDA detector (SPD-M40). LissRhod PE was detected with an excitation wavelength of 554 nm and emission of 576 nm. Each sample was injected at least twice to check reproducibility.

### Membrane protein expression

All tested membrane proteins were expressed in HEK293 suspension cells using the baculovirus expression system and a modified version of the BacMam vector pEG^69^. HEK293 cells were cultured at 37 °C, 8 % CO_2_. rSERCA1a (rabbit sarcoendoplasmic reticulum Ca^2+^ ATPase) with a C-terminal eGFP-StrepII tag and cASIC1 (chicken acid-sensing ion channel 1) with an N-terminal His_8_-eGFP tag were expressed in HEK293S GnTI^-^ cells (ATCC) and enhanced with 10 mM sodium butyrate (NaBu) 10 h post-transduction. Cells were harvested 72 h and 48 h post-transduction, respectively. hKCNK18 (human potassium channel subfamily K member 18) with an N-terminal StrepII-eGFP tag was expressed in FreeStyle^TM^ 293-F (HEK293F) cells (Thermo Fisher Scientific, TFS) and enhanced with 10 mM NaBu 10 h post transduction and harvested 48 h post-transduction. hTRPM4 (human transient receptor potential melastatin type 4) fused with an N-terminal StrepTagII-eGFP tag was expressed in HEK293F cells and enhanced with 10 mM NaBu 24 h post-transduction and collected 48 h post-transduction as described in Autzen *et al.*^45^.

### FSEC of membrane proteins in copolymer lipid nanodiscs

HEK293 cell pellets were resuspended in HBS supplemented with cOmplete^TM^ EDTA-free protease inhibitor cocktail (Sigma-Aldrich) at 50 µL/∼1 million cells/condition. 50 µL of the cell suspension was mixed with 50 µL 4% (w/v) polymer in HBS for a final concentration of 2% (w/v), briefly sonicated with a probe sonicator and incubated on a rolling table for 2-4 h at 4 °C. Large aggregates were removed from the suspension by ultracentrifugation at 113,700 xg for 15 min and the supernatant containing protein-filled lipid nanodiscs was filtered with a 0.22 µm spin filter before being loaded onto a Superose 6 Increase 5/150 GL column (Cytiva) equilibrated with HBS at 4 °C. The eluent was detected by a Shimadzu (Shimadzu Europa GmbH) liquid chromatography system equipped with an autosampler (SIL-40), a fluorometer (RF-20A), and a PDA detector (SPD-M40). eGFP was detected with an excitation wavelength of 488 nm and emission of 507 nm. Each sample was injected at least twice to check reproducibility.

### Purification of hTRPM4 in MAASTY_7.1_-50 lipid nanodiscs

Harvested hTRPM4-expressing HEK293F cell pellets were resuspended in HBS supplemented with Sigmafast^TM^ EDTA-free protease inhibitor cocktail (Sigma-Aldrich) and lyzed by pressure homogenizer (Avestin Emulsiflex) with 3 passes at 1500 psi. Lysates were centrifuged at 26,000 xg for 25 min (Sorvall RC 5B Plus) followed by membrane isolation through ultracentrifugation at 55,000 pm for 1 h (Beckman, Type 70 Ti Rotor). Membranes were resuspended in HBS by mortar and pestle at 20 mL/g and snap-frozen in LN_2_ before storage at −80 °C until further use. Purification of hTRPM4 was carried out similarly to the procedure described in Smith *et al.*^18^. In short, isolated membranes were solubilized in 2% (w/v) MAASTY_7.1_-50 (1 g/40 mL) for 2 h at 4 °C on a rolling table and insoluble fragments separated by centrifugation at 30,000 xg for 30 min. The supernatant was incubated overnight at 4 °C with StrepTactin beads (Cytiva) in HBS with 250 mM L-Arginine-HCl. Under gravity flow, the beads were washed in HBS with 250 mM L-Arginine-HCl, then the protein eluted upon addition of 5 mM desthiobiotin.

Protein-positive fractions were pooled and spin-concentrated to less than 500 µL. TEV protease was added at a 1:5 molar ratio (TEV:hTRPM4) and incubated overnight at 4 °C to remove the fusion tag. The sample was filtered with a 0.22 µm spin filter unit before loading onto a Superose 6 Increase 10/300 GL column (Cytiva) equilibrated with HBS on an AKTA Pure Protein Purification System (Cytiva). Peak fractions were pooled and spin-concentrated (determined by NanoDrop UV-vis spectrophotometer 280 nm, 1 mg = 1 Abs) before assessment by either single-particle cryo-EM, MALDI-TOF MS or lipidomics analysis. Eluted fractions from size-exclusion chromatography were assessed for purity by SDS-PAGE (Fig. S8).

### MALDI-TOF MS

Purified hTRPM4 in MAASTY_7.1_-50 nanodiscs (0.1 mL) were extracted with a 9:1 CHCl_3_/MeOH solution (0.3 mL). The extract was mixed with 2,5-dihydroxybenzoic acid (DHB) (20 mg/mL) in 9:1 CHCl_3_/MeOH matrix to a final concentration of 10 mg/mL of DHB. Mass spectra were recorded on a Bruker AutoflexTM MALDI-TOF MS Spectrometer.

### Lipidomics analysis

Purified hTRPM4 in MAASTY_7.1_-50 lipid nanodiscs (∼2 mg/mL) was mixed with trypsin (Promega, sequencing grade) in a 1:25 molar ratio in 100 mM ammonium bicarbonate pH 8, and digested overnight at 37 °C. Lyophilized mix of peptides and lipids was dissolved in 68% solution A (acetonitrile:H_2_O 60:40, 10 mM ammonium formate and 0.1% formic acid) and 32% solvent B (isopropanol:acetonitrile 90:10, 10 mM ammonium formate and 0.1% formic acid) for analysis by reverse-phase liquid chromatography tandem MS (RP LC–MS/MS). RP LC–MS/MS was performed using a Dionex UltiMate 3000 RSLC Nano system coupled to an Orbitrap Eclipse Tribrid Mass Spectrometer (TFS). The peptide/lipid mixture was loaded onto a C18 column (Acclaim PepMap 100, C18, 75 µm x 15 cm, Thermo Scientific^TM^) at a flow rate of 300 nL min^-1^. After 10 min, solvent B was ramped to 65% over 1 min, then 80% over 6 min, before being held at 80% for 10 min, then ramped to 99% over 6 min and held for 7 min for a total run of 45 min.

The mass spectrometer was set with a spray voltage of 2.4 kV and capillary temperature of 320 °C. Analysis was done in negative mode, as a data-dependant acquisition. Survey full-scan MS spectra were acquired on the Orbitrap (m/z 300–2,000) with a resolution of 120,000. Higher-energy Collision-induced Dissociation (HCD) fragmentation was done for the N most intense ions within 3 second cycle time, with max injection time of 50 ms and normalized collision energy of 25%, followed by a ramp to to 30%. The MS/MS analysis was done in the Orbitrap, at 15,000 resolution and the first mass at 75 m/z, max injection time of 50 ms and isolation window of 1.5 m/z.

Lipid identification was performed using LipiDex software (v.1.1) with the LipiDex HCD Formic library. The MS and MS/MS search tolerances were set to 0.01 m/z. List of potential adducts was manually curated, whereby a singly charged ion with formate adduct ([M + HCOO]^-^) was added to the list, as well as singly charged deprotonated ion ([M-H]^-^). Chromatogram integration and lipid quantification was carried out using MZmine (v.2.54), adapting the integrated LipiDex-MZmine workflow proposed by Hutchins *et al.*^70^.

### MP analysis

MP analyses were performed with a Refeyn Two^MP^ mass photometer (Refeyn Ltd). Data was acquired using AcquireMP (Refeyn Ltd, version 2.3). The instrument was calibrated using in-house prepared protein standards of Dynamin ΔPRD, which forms oligomeric configurations of the following masses: 90, 180, 360 kDa^71^. Strict linear response for calibration curves was observed and extrapolated with high confidence. Purified hTRPM4 in MAASTY_7.1_-50 lipid nanodiscs in its isolation buffer from SEC was diluted in phosphate-buffered saline (PBS) immediately before the MP acquisition to a final concentration of 50-300 nM, depending on the run. The focal plane was located using 15 *µ*L PBS, immediately after, the protein was mixed into the PBS droplet, diluting it 4 times in the process. The acquisitions were collected in the large field of view mode for 60 sec. The raw data were processed and manually curated using DiscoverMP (Refeyn Ltd, version 2.3) to calculate the histograms of the binding events for mass determination and integrate them for quantification of different mass species. Factory-set video processing settings were used for mass histogram calculations.

### Determination of the hTRPM4-MAASTY nanodisc composition

Harvested HEK293F cell membranes expressing hTRPM4 were solubilized with 1% (w/v) MAASTY_7.1_-50_ttc_ and purified as described above. Final protein concentration was determined through Bradford assay against a BSA standard curve. A calculated 10 µg purified hTRPM4 was injected onto a Superose 6 Increase 5/150 gel filtration column and measured for absorbance at 280 nm (protein) and 310 nm (ttc). Specified concentrations of MAASTY_7.1_-50_ttc_ copolymer were injected onto a Superose 6 Increase 5/150 gel filtration column and measured at 310 nm absorbance to construct a standard curve of ttc concentration. Total MAASTY_7.1_-50_ttc_ per mole hTRPM4 was then calculated using the peak 310 nm absorbance values. Remaining lipid content was calculated using the MP analysis, which determined the molecular weight per particle, the molecular weight of hTRPM4, and the molecular weight of the encasing copolymers.

### Purification of hTRPM4 in MSP2N2 lipid nanodiscs

hTRPM4 was purified in n-Dodecyl-beta-D-maltoside (DDM) and cholesteryl hemisuccinate (CHS) and reconstituted in MSP2N2 nanodiscs as described in Autzen *et al.*^45^. The final concentration of the sample was 2.2 mg/mL to which CaCl_2_ was added to a final of 5 mM just before vitrification.

### Negative stain EM of hTRPM4 and HEK293F membranes

EM grids of negatively stained hTRPM4-MAASTY nanodiscs or MAASTY solubilized HEK293 membranes were prepared by applying 3 µL of sample (0.06-0.3 mg/mL for hTRPM4-MAASTY, determined by Implen UV-vis NanoPhotometer 280 nm, 1 mg = 1 Abs) to a 400-mesh copper grid covered with continuous carbon film and stained with 2 % (w/v) uranyl acetate. Negative-stain EM grids were imaged on a CM100 microscope (Philips) operated at 100 kV and equipped with an Olympus Veleta camera. Images were recorded at a nominal magnification of 135,000x, corresponding to a pixel size of 3.8 Å.

### Cryo-EM sample preparation and data collection

Cryo-EM grids were prepared by adding 4 µL of ∼1.4 mg/mL purified hTRPM4-MAASTY or 2.2 mg/mL hTRPM4-MSP2N2 in lipid nanodiscs to glow-discharged Quantifoil R1.2/1.3 300-mesh Au holey carbon grids (Quantifoil, Micro Tools GmbH). The grids were blotted with Whatman Grade 1 filter paper for 3 s at 4 °C and 100% humidity, then plunge-frozen into liquid ethane using a by Vitrobot^TM^ Mark IV (TFS). Datasets were collected on a Titan Krios G2cA cryo-transmission electron microscope equipped with a Thermo Scientific^TM^ Selectris X imaging filter and a Falcon4i detector. Grid squares were manually selected and movies were recorded in counting mode using EPU (TFS) at a nominal magnification of 165kx, corresponding to a physical pixel size of 0.725 Å. Data were collected with a defocus range of 0.8 to 2.5 µm and a total electron dose of 52 e^-^/Å^2^. Images were saved as Electron Event Recordings (EER). Details on data collections are included in Table S1.

### Image processing

Pre-processing of both nanodisc samples was performed in cryoSPARC live v4.4.0 for initial assessment of sample quality during which patch-based motion correction was used for aligning EM movie stacks, and applying dose-dependent weighting^72^. Further processing was performed using a combination of cryoSPARC v4.4.0^72^ and Relion 5.0^73^.

For hTRPM4 in MAASTY_7.1_-50 nanodiscs, a total of 6308 movies at 0.749 Å/pixel were collected. This predetermined pixel size was later corrected to 0.725 Å/pixel in post-processing following an accurate determination of pixel size, better reflecting the detector specifications. In cryoSPARC v4.4.0, motion corrected micrographs were patch-based CTF corrected before curating to 6070 micrographs. Particle picking was first carried out using blob picker, from which two rounds of 2D classifications identified classes used as templates for template-based picking. These 1,306,897 particles underwent three rounds of 2D classification leaving 126,417 particles remaining. An *ab initio* model was generated and classified using heterogeneous refinement. The best class of 87,697 particles was refined further by non-uniform refinement, resulting in a reconstruction with an estimated resolution of 3.23 Å. Due to lower local resolution in the cytosolic regions, we re-extracted the refined particles in Relion 5.0, after particle stack conversion by pyEM^74^, from micrographs motion-corrected using the CPU-based implementation of MotionCor2^75^. The particles underwent refinement applying C4 symmetry and BLUSH regularization^76^ resulting in a 3.46 Å resolution reconstruction following post-processing with a soft mask. Details on image processing and final reconstructions of hTRPM4 in MAASTY nanodiscs are included in Fig. S10, S11, and Table S1.

For this study, we solved the hTRPM4-MSP2N2 structure from cryo-EM data collected on gold grids in contrast to the structures from Autzen *et al.*^45^ which were solved from data collected on copper grids. For hTRPM4 in MSP2N2 nanodiscs, a total of 7,945 EER format movies at 0.725 Å/pixel were collected and imported into Relion 5.0^73^. Movies were motion corrected with the CPU-based implementation of MotionCor2 using an EER fractionation of 40, and dividing whole frames into 5 × 5 patches^75^.

Micrographs were CTF corrected with ctffind 4.1^77^, manually curated and micrographs with resolution lower than 6 Å were discarded, resulting in 7627 micrographs to pick particles from. Particle picking was first carried out using a Laplacian-of-Gaussian (LoG) filter from which a round of 2D classifications helped identify classes that were used as templates for template-based picking. 1,104,004 particles were subjected to a single round of 2D classification (EM algorithm, 25 iterations) resulting in 255,586 good looking particles. These particles were used for determination of an *ab initio* model and subjected to 3D classification. The particles belonging to the two best looking classes were combined and refined in 3D with BLUSH regularization^76^ resulting in a reconstruction with an estimated resolution of 3.8 Å. Details on image processing and final reconstructions of hTRPM4 in MSP2N2 nanodiscs are included in Fig. S13, Fig. S12 and Table S1.

### Model building and refinement

Atomic models for hTRPM4 in MAASTY_7.1_-50 and MSP2N2 were built using Coot v0.9.8.94^78^ using the hTRPM4-MSP2N2 Ca^2+^ model 6BQV^45^ as the initial model. Residues missing from the reconstructions were not modeled. In the case of the MAASTY_7.1_-50 structure, this included MHR1-3, in addition to residues predicted to be unstructured which were missing from both structures. Table S1 provides an overview of the residues included in the final, refined models in addition to their refinement and validation statistics by Phenix and MolProbity (within Phenix version 1.21.1-5286)^79,80^. Both models were subjected to real-space refinement using Phenix v1.21.1-5286 with a general restaints setup. As cross-validations, the final models were refined against the finalized maps generated by Relion 5.0 (Fig. S10; Fig. S13). The program HOLE^81^ was used to calculate the pore profiles shown in Fig.3e. UCSF ChimeraX was used to prepare figures^82^.

## Data Availability

The reconstructed maps are available from the EMDB under access codes EMD-52611 (hTRPM4 in MAASTY) and EMD-????? (hTRPM4 in MSP2N2). The atomic models are available from the PDB under access codes PDB 9I3R (hTRPM4 in MAASTY) and(hTRPM4 in MSP2N2). The raw cryo-EM movies data used in our study are available under access codes EMPIAR-????? and EMPIAR-?????. Plasmids and other data that support the findings of this study are available from the corresponding authors upon reasonable request. Source data are provided in this paper.

## Code Availability

The code used for data analysis in this study was implemented using Python 3.10.6. All scripts and computational workflows necessary to reproduce the results are available upon reasonable request from the corresponding authors.

## Acknowledgements

The authors thank senior research scientists Tillmann Hanns Pape, Nicholas Heelund Sofos and Michael James Johnson for their support for cryo-EM experiments at the Core Facility for Integrated Microscopy at University of Copenhagen. The authors thank members of the Autzen Group for data discussion. hTRPM4 was expressed in a modified version of a BacMam vector gifted from Eric Gouaux (Addgene plasmid # 160680). Subsets of the figures were created with BioRender.com.

## Author Contributions

H.E.A., and A.A.A.A. conceived the project. C.F.P., H.E.A., and A.A.A.A. designed the experimental procedures. A.A.A.A., K.F.P., C.S., and N.J.B. carried out copolymer synthesis. N.J.B. carried out NMR. C.F.P. carried out expression of all membrane proteins, FSEC screening of membrane proteins and fluorescent lipid nanodiscs, purification and vitrification of hTRPM4 in MAASTY nanodiscs, and preparation of hTRPM4 for lipidomics and MP analysis. L.P.F. purified hTRPM4 in MSP2N2 nanodiscs and C.d.L vitrified the sample for cryo-EM. C.F.P. and L.P.F. performed image acquisition. C.F.P. and H.E.A. performed single particle analysis. C.F.P. built atomic models. K.F.P. carried out calcium binding experiments and pH sensitivity. A.A.A.A. did MALDI-TOF analysis of hTRPM4 in MAASTY. D.Z. and J.R.B. carried out lipidomics analysis and MP. C.F.P., H.E.A. and A.A.A.A wrote the manuscript. C.F.P., H.E.A. and A.A.A.A. prepared figures. All authors read and reviewed the manuscript.

## Inclusion and Ethics

In preparing this manuscript, we ensured equitable recognition of all contributors. This study did not involve human participants, human tissues, or animal research requiring ethical approval. The research adheres to Nature Portfolio’s policies on integrity, accessibility, and reproducibility, ensuring transparency and rigor in scientific reporting.

## Supplementary Information

The accompanying supplementary figures and tables are provided as a separate file.

## Supporting Information

**Figure S1:**
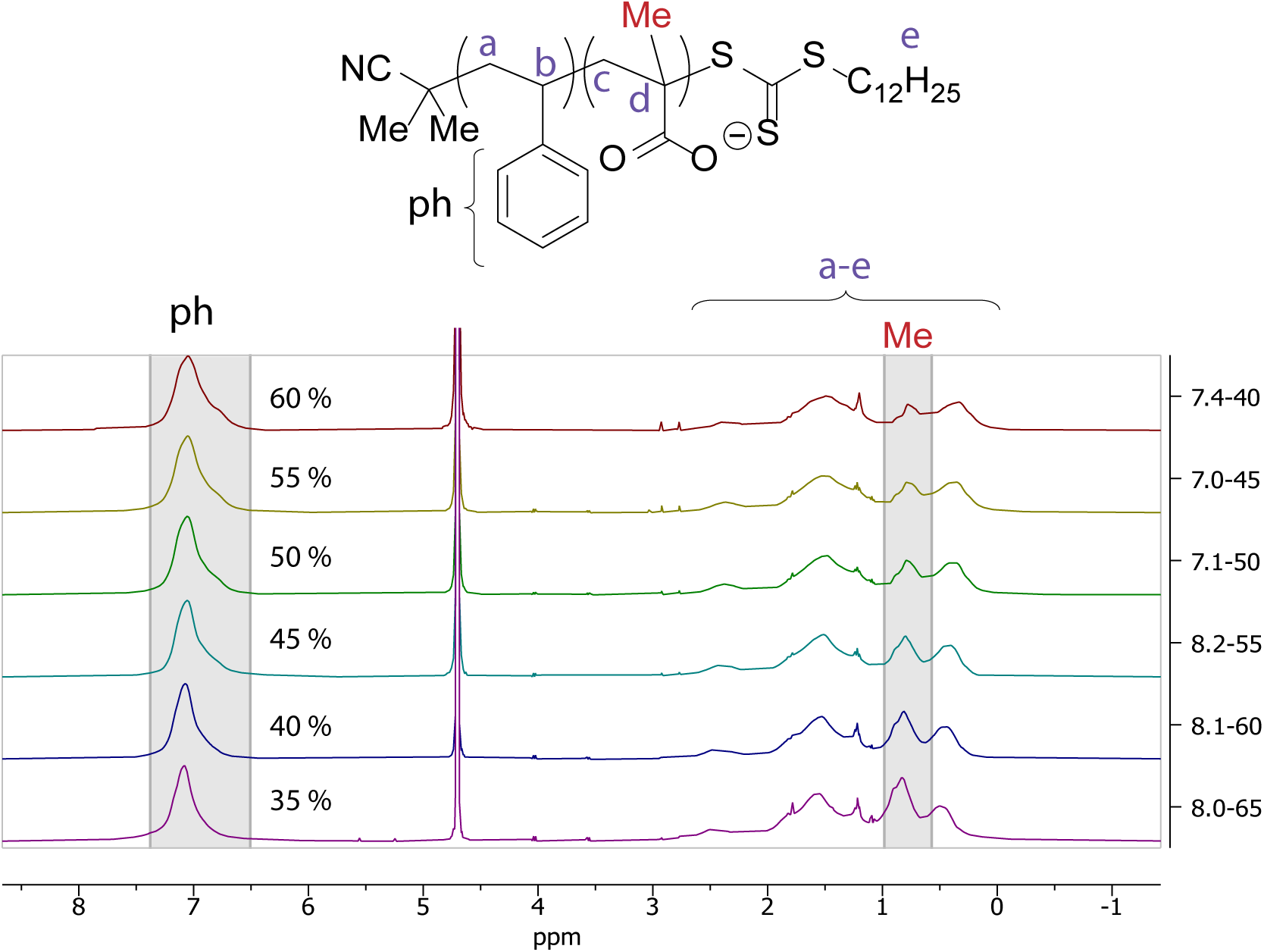
^1^H-NMR of MAASTY copolymers. ^1^H-NMR (D_2_O, 400 MHz): *δ* [ppm] = 7.5 - 6.5 (phenyl H), 3.0 - 0.3 (aliphatic H). STY content is seen in the part of the spectrum marked by ph (shaded in grey, curly brackets in structure). The shoulder at 6.8 ppm arises from an increase in adjacent STY-STY monomers with increasing STY content, which changes the local chemical environment for the protons in the phenyl. The methyl signal from MAA is seen to increase in proportion with MAA content in the part of the spectrum marked by Me (shaded in grey, red in structure). The MAA content is in correspondence with the initial feed of MAA.

**Figure S2:**
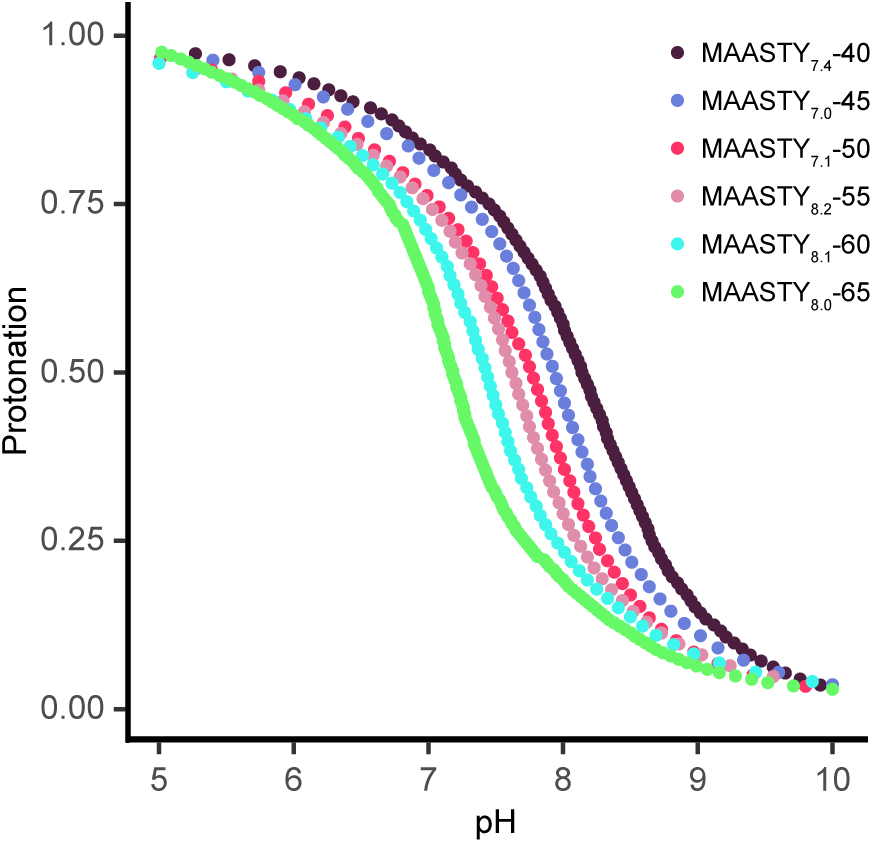
Bjerrum diagram of MAASTY copolymers describing copolymer pKa and pH tolerance. Bjerrum diagrams were constructed by dissolving 50 mg of each MAASTY copolymer in 10 mL water and adjusting the pH to approximately 11 with 1 M NaOH. The pH was measured on a Mettler Toledo SevenCompact S210 pH meter while 0.33 M HCl solution was titrated in 5 µL increments into the dissolved copolymer solution.

**Figure S3:**
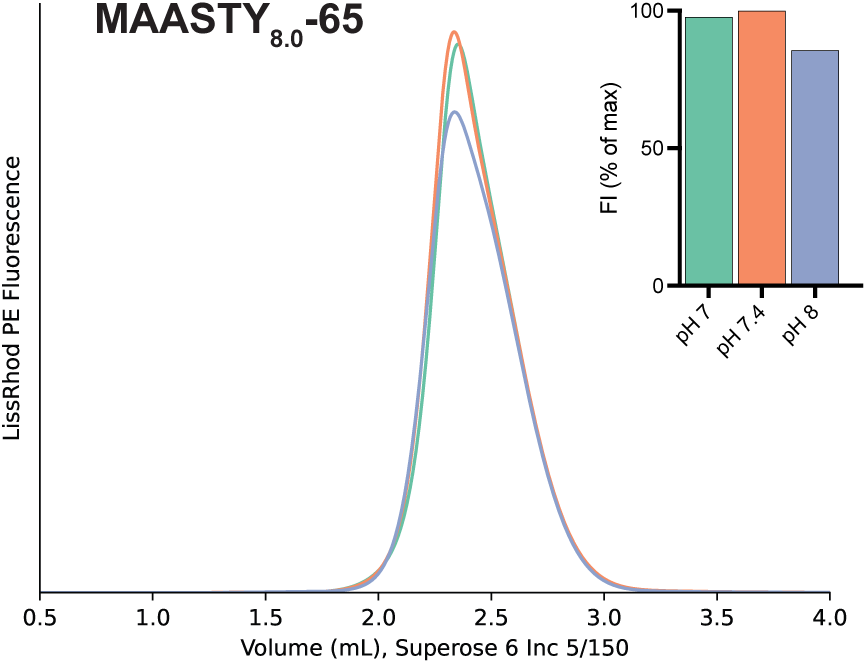
MAASTY_8.0_-65 solubilization of lipid vesicles at different pH. Fluorescent, protein-free nanodiscs were produced as described in methods and as illustrated in Fig. 5a. Raw FSEC traces of protein-free nanodiscs generated by solubilizing 1 mM POPC with 2% fluorescent LissRhod PE lipid vesicles with 1% (w/v) of MAASTY_8.0_-65 in specified pH. Samples were injected onto a Superose 6 Increase 5/150 gel filtration column and detected for LissRhod PE with excitation at 554 nm and emission at 576 nm. Insert bar graph represents maximum fluorescence intensities of the traces, normalized to maximum detected peak intensity. Each bar represents a single data point.

**Figure S4:**
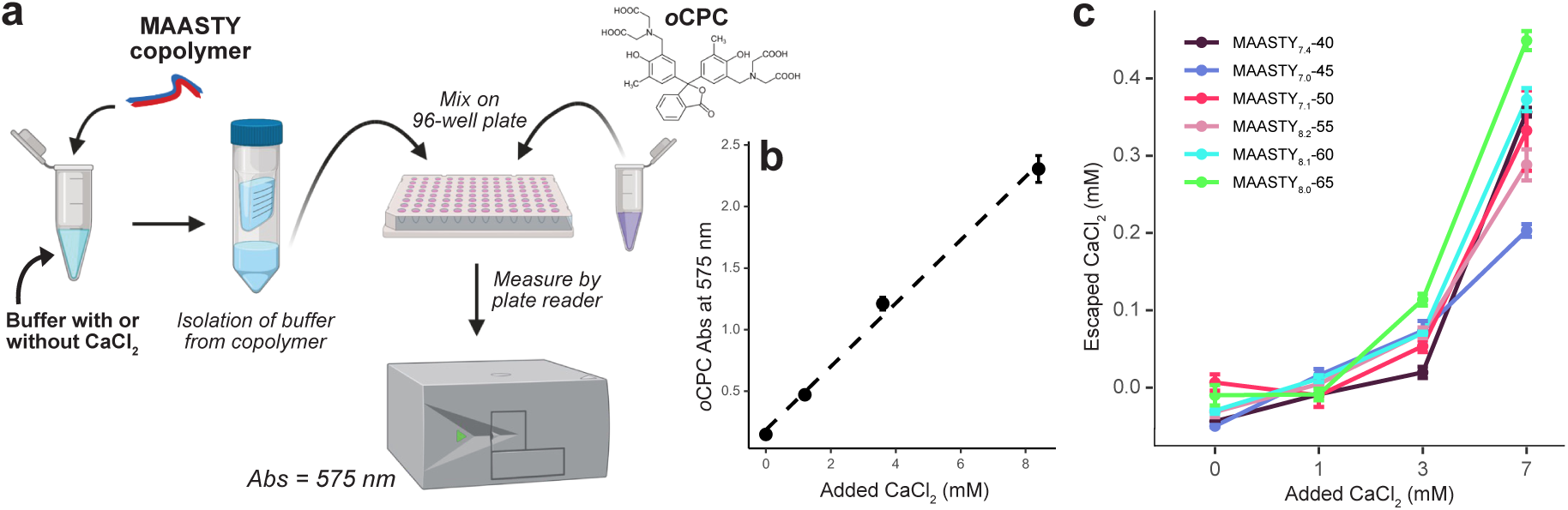
**Assessment of Ca**^2+^ **sensitivity of MAASTY copolymers with o-cresolphthalein complexone (oCPC). a** Schematic representation of the assay. Tested MAASTY copolymers were solubilized and mixed in buffers of examined CaCl_2_ concentrations. Buffer was then isolated from the copolymer using a 10 kDa MWCO centrifugal concentrator and mixed with oCPC in a 96-well plate. Finally, absorbance at 575 nm was measured in a plate reader to calculate free Ca^2+^ concentration. **b** Standard curve of measured oCPC in increasing CaCl_2_ concentrations. Data points represent triplicate measurements ± standard deviation (SD, error bars), R^2^ = 0.995. **c** Free Ca^2+^ concentration in solution following exposure to 1% MAASTY copolymer as measured using oCPC. Data points represent triplicate measurements ± SD (error bars).

**Figure S5:**
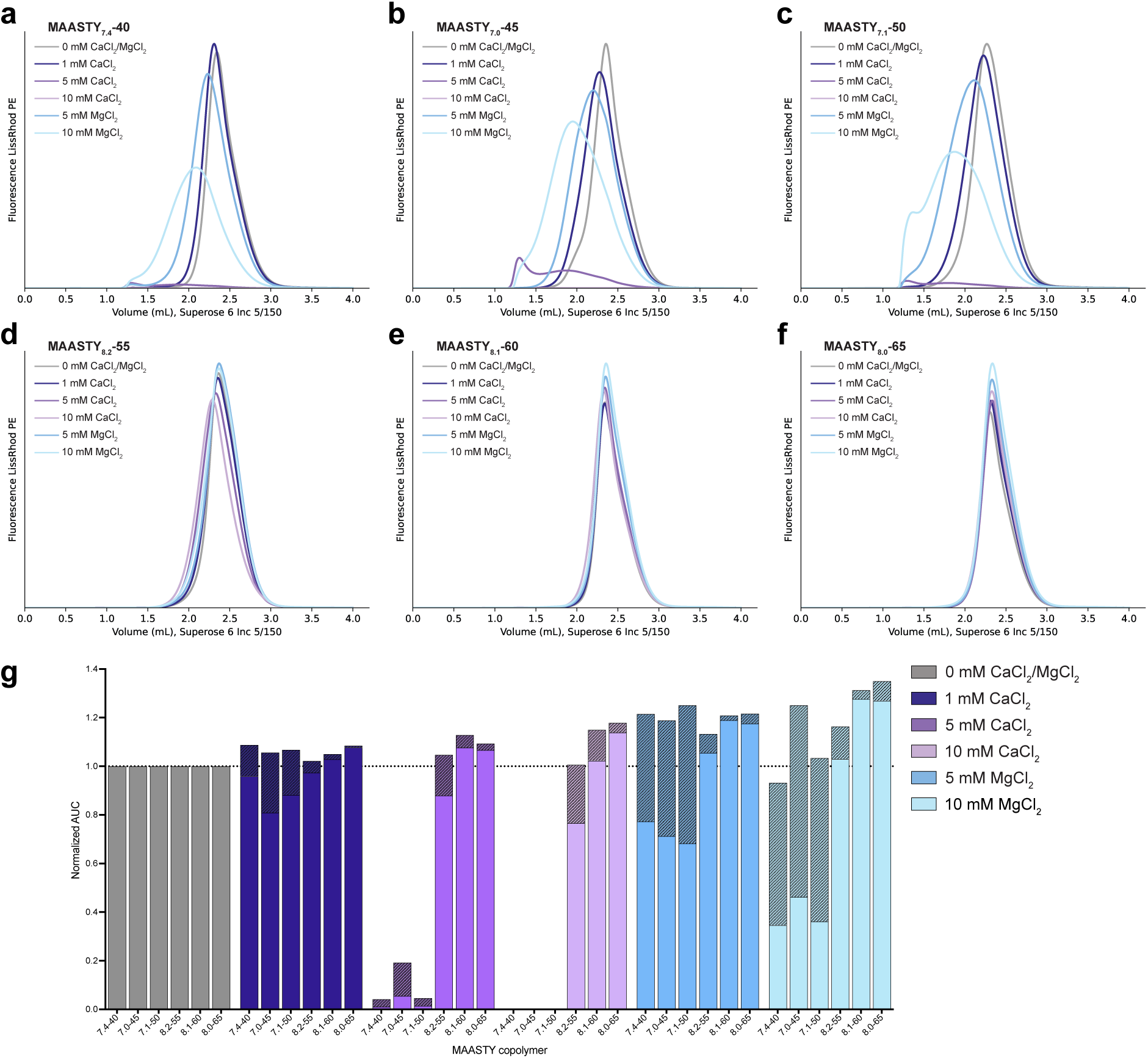
Solubilization of lipid vesicles by MAASTY copolymers in the presence of divalent cations. Fluorescent, protein-free nanodiscs were produced as described in methods and as illustrated in Fig. 5a. **a-f** Raw FSEC traces of protein-free nanodiscs generated by solubilizing 1 mM POPC with 2% fluorescent LissRhod PE lipid vesicles with 1% (w/v) of a given MAASTY copolymer in the presence of specified CaCl_2_ and MgCl_2_ concentrations. Samples were injected onto a Superose 6 Increase 5/150 gel filtration column and detected for LissRhod PE with excitation at 554 nm and emission at 576 nm. **g** Analysis of area under the curve (AUC) for the nanodisc peak which elutes at a retention volume of ∼2.3 mL. Each datapoint is normalized to the AUC of the 0 mM CaCl_2_/MgCl_2_ condition for each respective MAASTY copolymer. The nanodisc-containing peak for each condition was determined as that which overlaps with the 0 mM divalent cation condition (solid color). Bar sections with a diagonal line fill represent the larger, soluble species determined as AUC from a lower retention volume than the 0 mM divalent cation condition. Each bar represents a single datapoint.

**Figure S6:**
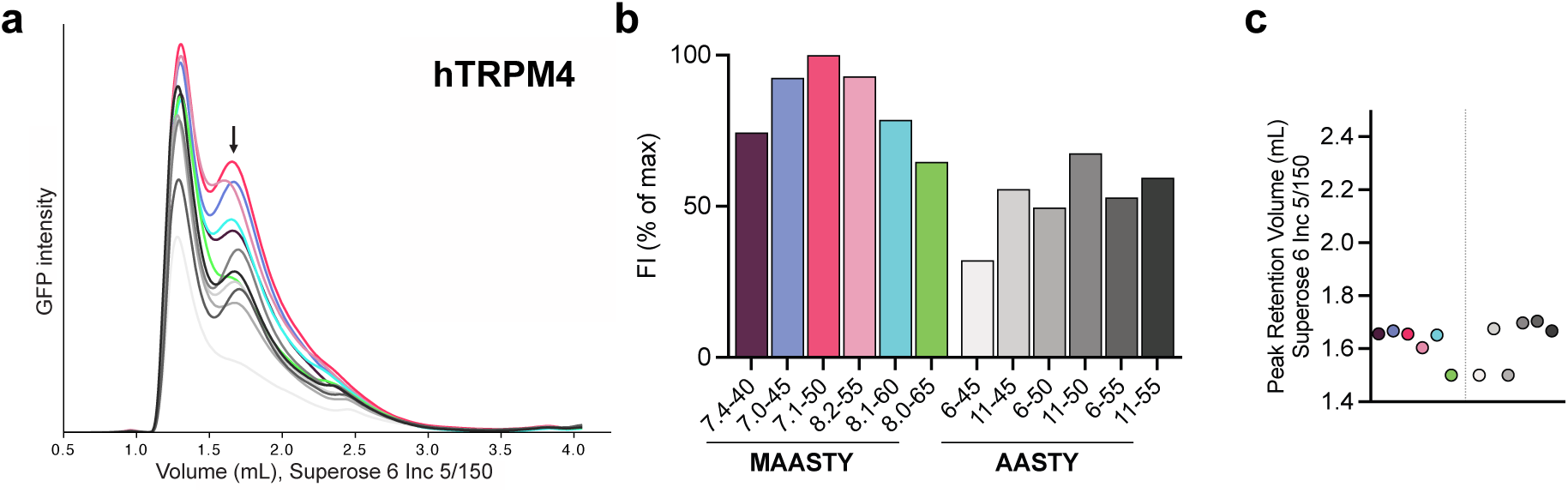
Solubilization of hTRPM4 in MAASTY copolymers. **a** Raw FSEC traces of hTRPM4 solubilized by the MAASTY library presented in this study and for comparison the AASTY library presented in Smith *et al.*^1^. The samples were injected onto a Superose 6 Increase 5/150 gel filtration column and detected for eGFP using an excitation wavelength of 488 nm and emission of 507 nm. From the chromatogram, a void peak is seen at approximately 1.2 mL with the presumed protein-nanodisc peak denoted by an arrow. **b** Maximum fluorescence intensities of the traces in **a**, normalized to maximum detected peak intensity. **c** Peak retention volumes of the traces in **a**. Each unique membrane protein copolymer sample was injected twice or more to check reproducibility.

**Figure S7:**
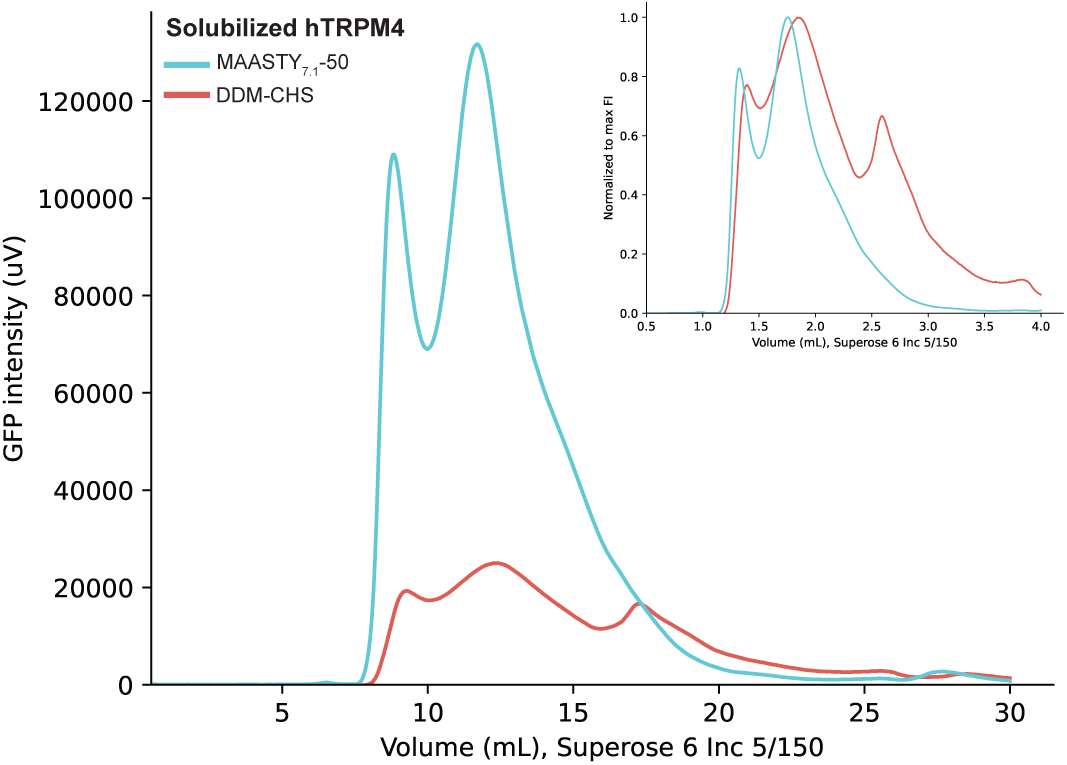
Solubilization comparison of hTRPM4 by MAASTY_7.1_-50 and DDM-CHS. Raw FSEC traces of hTRPM4 solubilized by 2% (w/v) MAASTY_7.1_-50 (blue) and 20 mM DDM with 4 mM CHS (red). The samples were injected onto a Superose 6 Increase 5/150 gel filtration column equilibrated in HBS or buffer containing 100 mM NaCl, 0.5 mM DDM and 20 mM HEPES/NaOH, pH 7.4 for the MAASTY or DDM-CHS conditions, respectively, and detected for eGFP using an excitation wavelength of 488 nm and emission of 507 nm. Insert represents each trace normalized to maximum GFP intensity showing the distribution of each trace.

**Figure S8:**
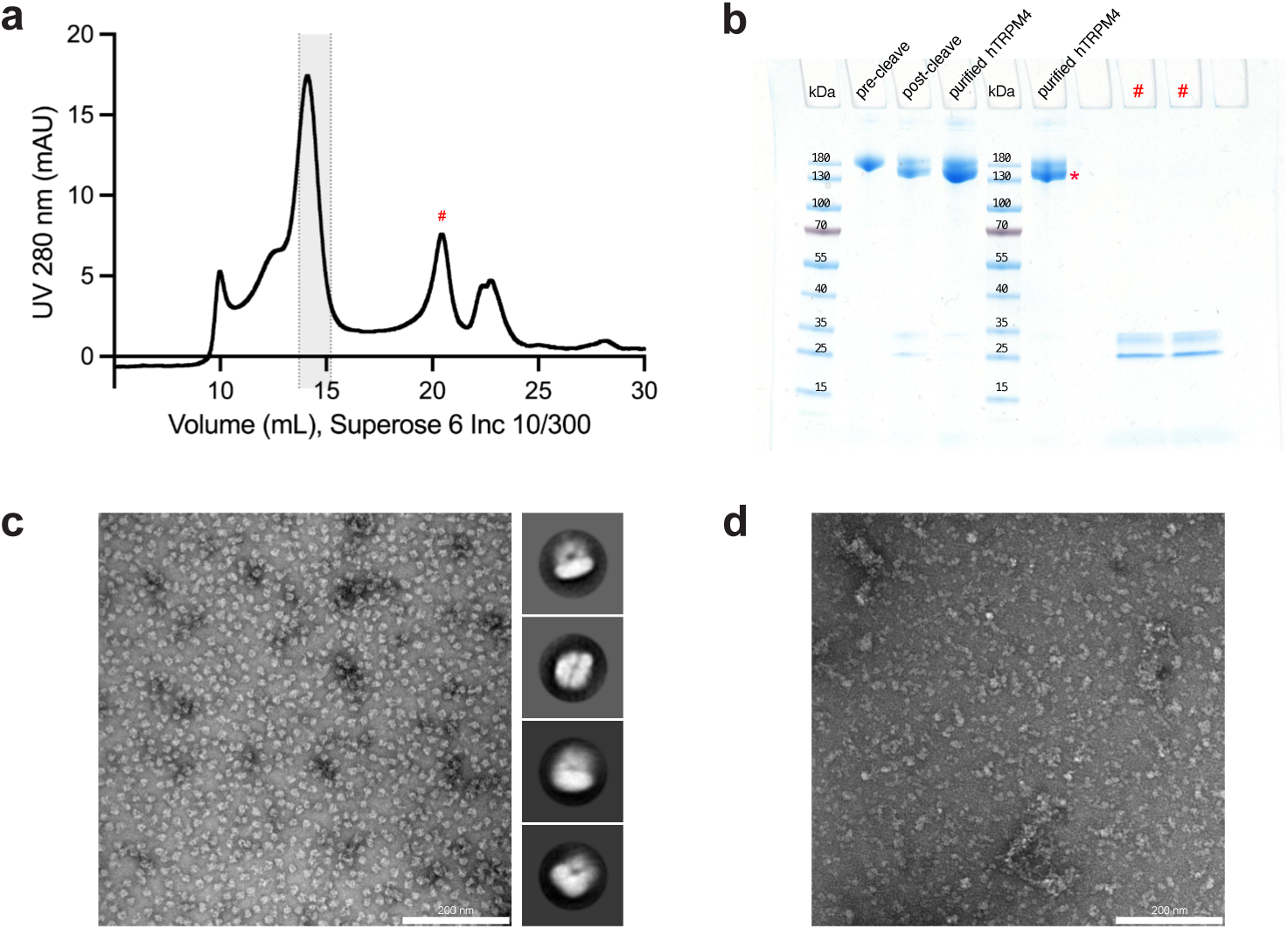
Purification of hTRPM4 in MAASTY lipid nanodiscs. **a** Chromatogram of absorbance at 280 nm of purified hTRPM4 in MAASTY native nanodiscs from Superose 6 Increase 10/300 gel filtration column. Sample fractions (highlighted in grey) were pooled and concentrated for structural determination of hTRPM4 in MAASTY by cryo-EM. The peak at 10 mL and shoulder at ∼12.5 mL derive from the void and heterogeneous aggregates of hTRPM4, respectively. The peak at 20 mL contains TEV protease and the StrepTagII-eGFP tag as analyzed by SDS-PAGE in **b** in the lanes denoted by #. **b** SDS-PAGE analysis of hTRPM4-containing MAASTY native nanodiscs representing (from left to right): hTRPM4-MAASTY before the addition of TEV protease, hTRPM4-MAASTY after O/N cleavage by TEV protease, purified hTRPM4 in MAASTY native nanodiscs for downstream analyses (repeated), 2 fractions from the peak denoted by # in **a**. The protein band at ∼134 kDa represents the proteomer of hTRPM4 in MAASTY nanodiscs and is marked by *. The faint band above is partially uncleaved purified hTRPM4 in MAASTY as identified by in-gel fluorescence (data not shown).**c** Negative stain TEM image of hTRPM4-MAASTY with representative 2D classes. **d** Negative stain TEM image of MAASTY-solubilized HEK293F membranes expressing hTRPM4.

**Figure S9:**
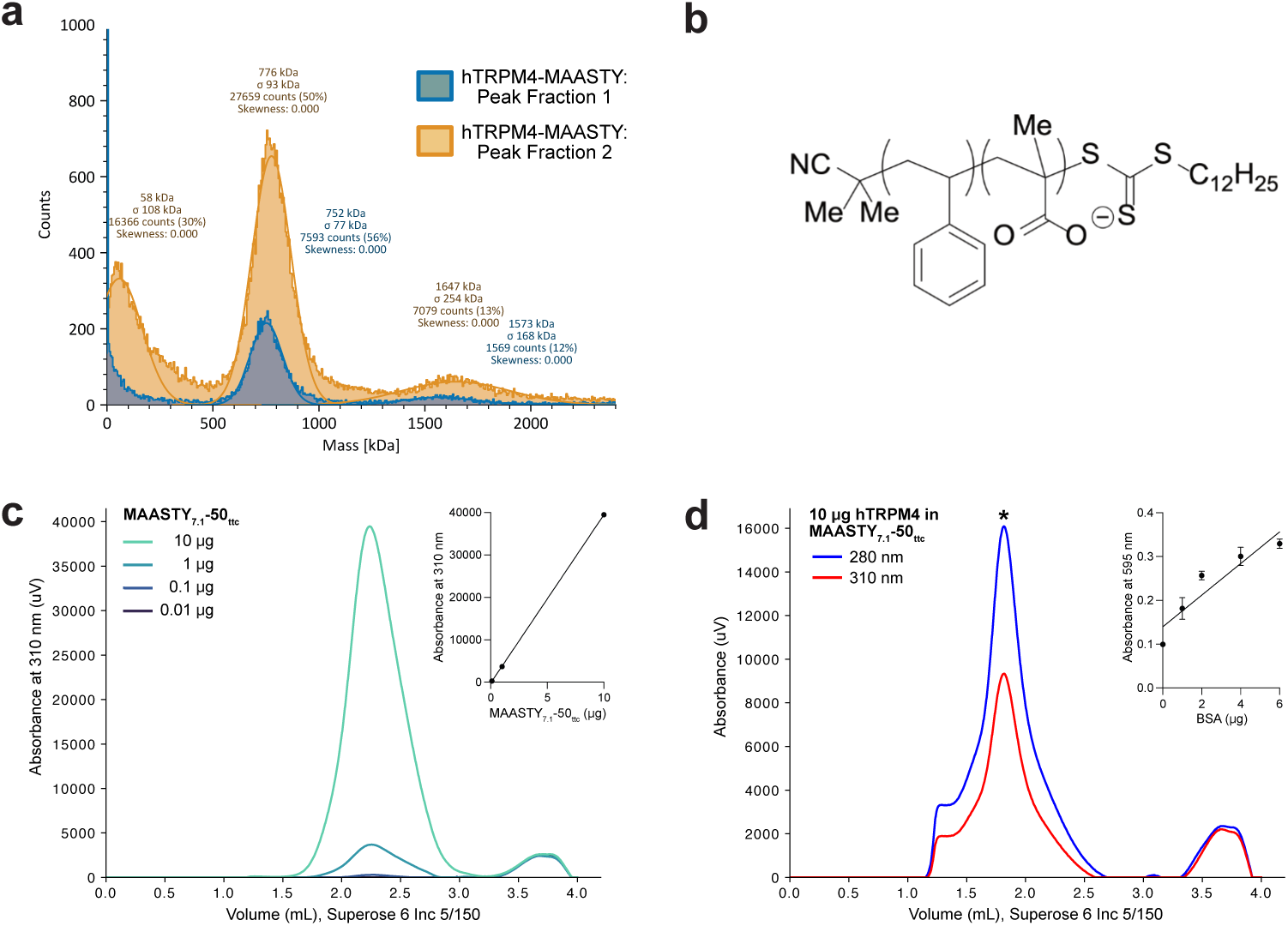
Compositional analysis of hTRPM4 in MAASTY lipid nanodiscs. **a** Mass photometry analysis of peak fractions from purification of hTRPM4 in MAASTY nanodiscs as shown in Fig. S8a. Two 500 *µ*L fractions of purified hTRPM4-MAASTY were collected and analyzed. Peak fraction 1 (blue) comprises the majority of the protein peak with hTRPM4-MAASTY likely representing the 760 kDa species, whereas peak fraction 2 (orange) comprises the right-side of the peak (larger retention volume) as evident by the higher abundance of smaller species. **b** Molecular structure of MAASTY copolymer with the trithiocarbonate (ttc) moeity (RAFT reagent), which has a maximum absorbance of 310 nm. **c** Raw SEC traces of specified masses of MAASTY_7.1_-50_ttc_ copolymer injected on to a Superose 6 Increase 5/150 gel filtration column and measured at 310 nm absorbance. Insert represents a standard curve derived from the maximum intensity values from the copolymer peak (∼2.3 mL retention volume). **d** Raw SEC traces from the injection of 10 *µ*g purified hTRPM4 in MAASTY_7.1_-50_ttc_ lipid nanodiscs on to a Superose 6 Increase 5/150 gel filtration column measured at 280 nm (blue) and 310 (red) nm absorbance, representing total protein and MAASTY_7.1_-50_ttc_ copolymer, respectively. Insert represents a BSA standard curve from the Bradford protein assay used to determine the concentration of purified hTRPM4 in MAASTY_7.1_-50_ttc_ lipid nanodiscs prior to SEC.

**Figure S10:**
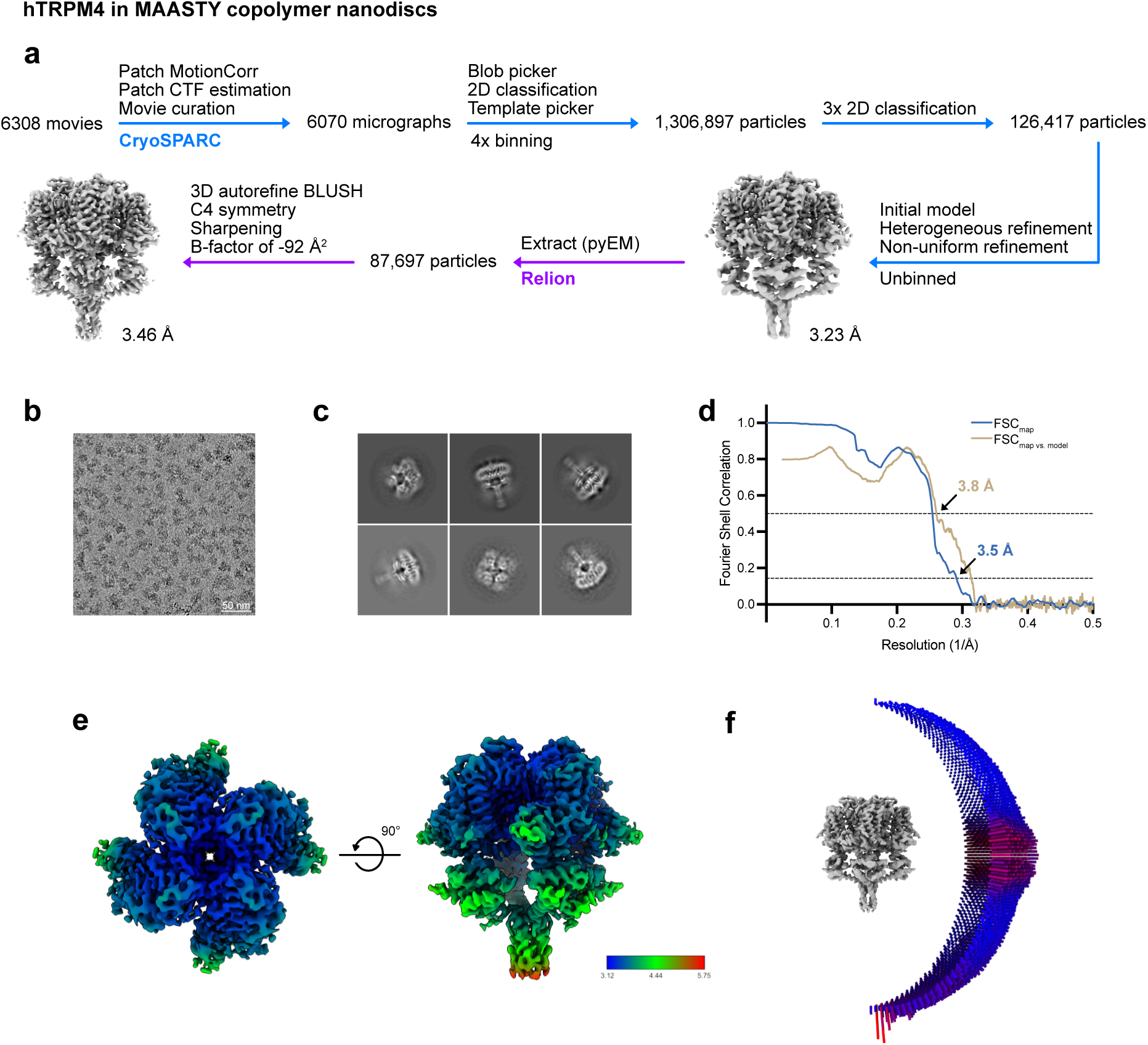
Cryo-EM data processing of hTRPM4 in MAASTY lipid nanodiscs. **a** Cryo-EM data processing pipeline for hTRPM4 in MAASTY. Motion correction, CTF estimation, particle picking, 2D classification and primary refinement steps were carried out in CryoSPARC^2^ (blue arrows). Refined particles were then re-extracted in Relion 5.0^3^ after particle stack conversion by pyEM^4^, and refined using BLUSH regularization^5^ (purple arrows). Despite an overall slightly lower estimated resolution, the reconstructions obtained from refinement in Relion 5.0 with BLUSH regularization and sharpening showed higher occupancy and better-resolved cytosolic domains than the map determined from non-uniform refinement in CryoSPARC. This led us to continue with the Relion 5.0 reconstructions, with an estimated overall resolution of 3.46 °A, for further analyses. **b** Representative motion-corrected micrograph imaged at 165,000x magnification. **c** Representative 2D class averages. **d** Fourier shell correlation (FSC) half-map and model-map curves, adopted from Relion 5.0 and Phenix Mtriage^6^, respectively. The gold standard FSC curve from the two half maps (FSC_map_) is shown in blue with resolution at FSC = 0.143 indicated by an arrow. The FSC curve generated from the map and atomic model (FSC_map vs. model_) is shown in brown with resolution at FSC = 0.5 indicated by an arrow. **e** Local resolution representation of the best reconstruction. **f** Angular distribution of particles used in the final reconstruction.

**Figure S11:**
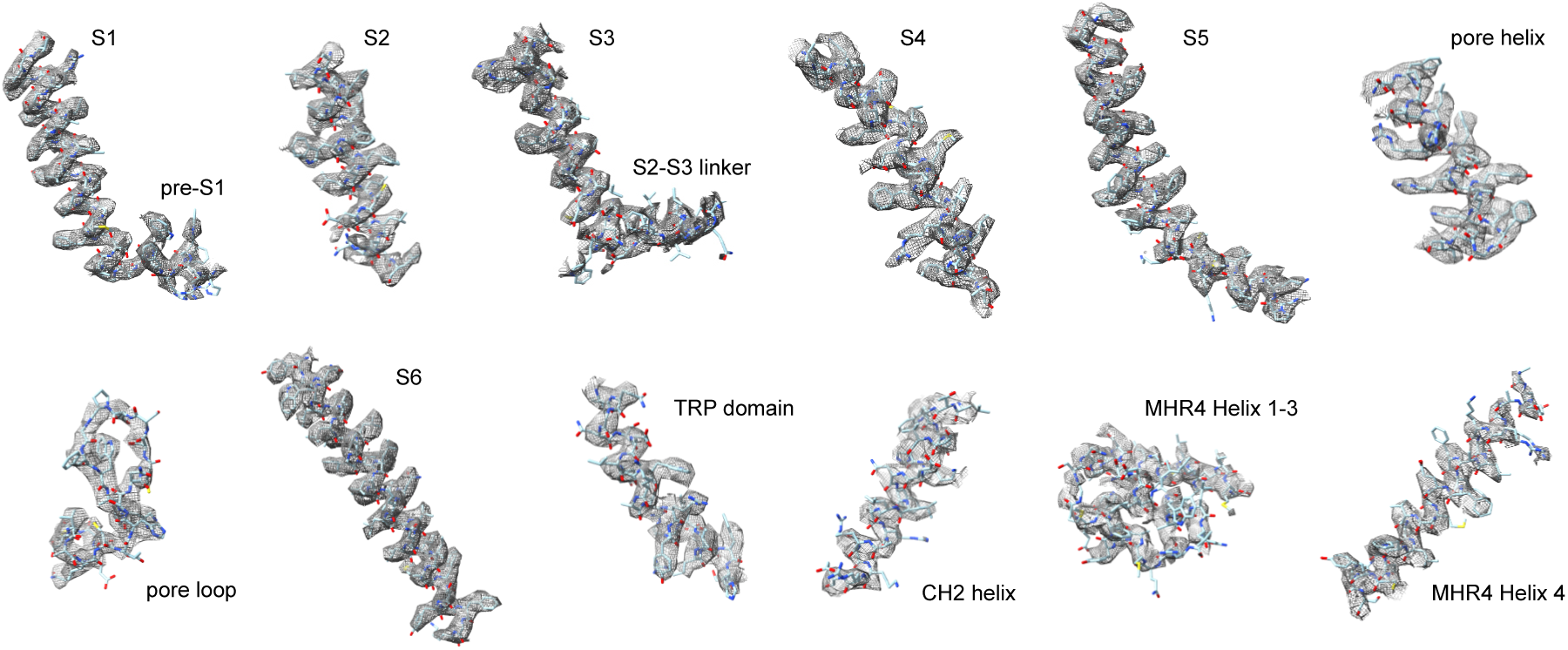
Representative regions from the best reconstruction of hTRPM4 in MAASTY nanodiscs. Shown are representative labelled segments of hTRPM4 with the fitted atomic model.

**Figure S12:**
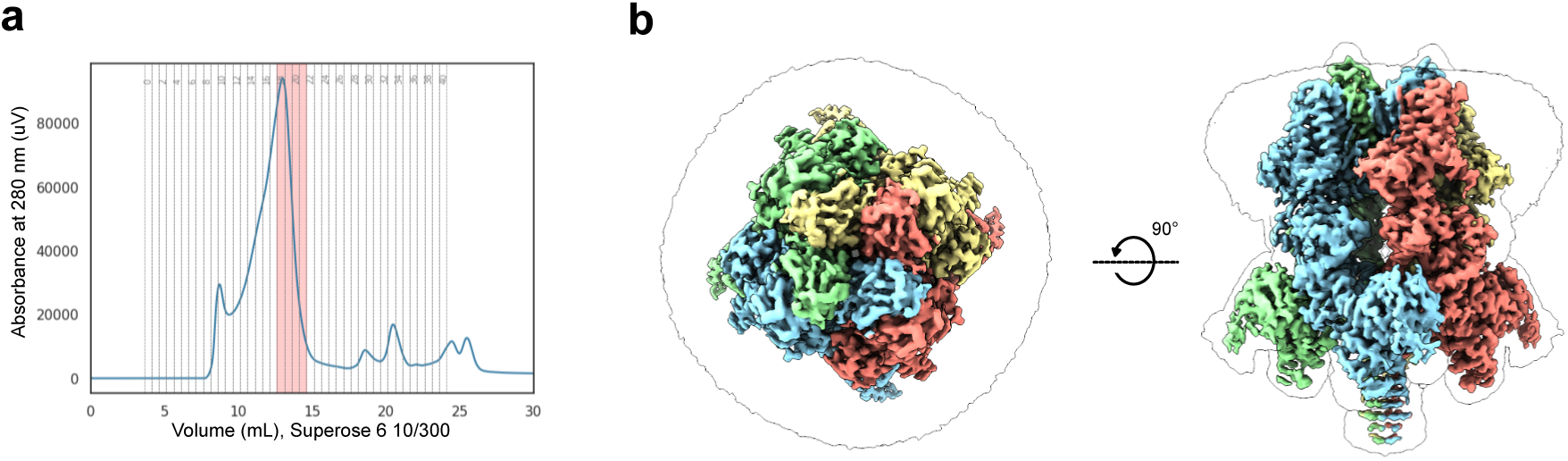
Purification and structural determination of hTRPM4 in MSP2N2 reconstituted nanodiscs. **a** Chromatogram of absorbance at 280 nm of purified hTRPM4 in MSP2N2 nanodiscs from Superose 6 Increase 10/300 gel filtration column. Sample fractions (highlighted in red) were pooled and concentrated for structural determination of hTRPM4 in MSP2N2 by cryo-EM. **b** Side and top views of sharpened cryo-EM density maps of hTRPM4 in MSP2N2 reconstituted nanodiscs with the four protein subunits colored in light green, salmon, sky blue and khaki, overlaid with the unsharpened cryo-EM density map (black outline). The unsharpened map represents the raw reconstruction, showing the full range of densities, including weaker and flexible regions such as copolymers and lipids in the nanodiscs. The sharpened map has been processed with a B-factor of −147 °A² to enhance high-resolution features, improving clarity in well-ordered regions. Contour levels for both maps were adjusted for optimal visualization of key structural features.

**Figure S13:**
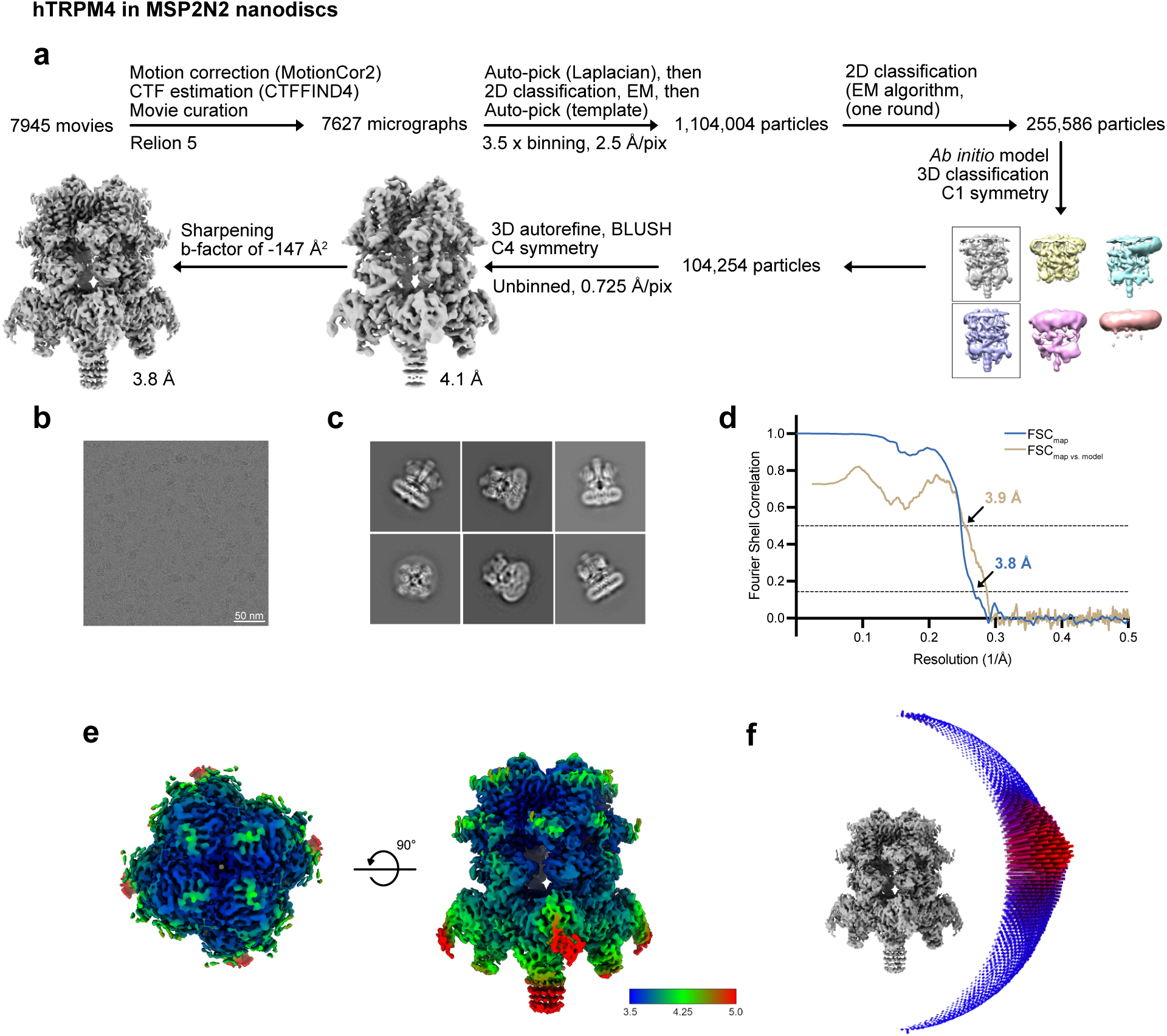
Cryo-EM data processing of hTRPM4 reconstituted in MSP2N2 lipid nanodiscs. **a** Cryo-EM data processing pipeline for hTRPM4 in MSP2N2. Motion correction, CTF estimation, particle picking, 2D classification, 3D classification and 3D refinement were carried out in Relion 5.0^3^. Blush regularization was applied during 3D refinement^5^. **b** Representative motion-corrected micrograph imaged at 165,000x magnification. **c** Representative 2D class averages. **d** Fourier shell correlation (FSC) half-map and model-map curves, adopted from Relion 5.0 and Phenix Mtriage^6^, respectively. The gold standard FSC curve from the two half maps (FSC_map_) is shown in blue with resolution at FSC = 0.143 indicated by an arrow. The FSC curve generated from the map and atomic model (FSC_map vs. model_) is shown in brown with resolution at FSC = 0.5 indicated by an arrow. **e** Local resolution representation of the best reconstruction. **f** Angular distribution of particles used in the final reconstruction.

**Figure S14:**
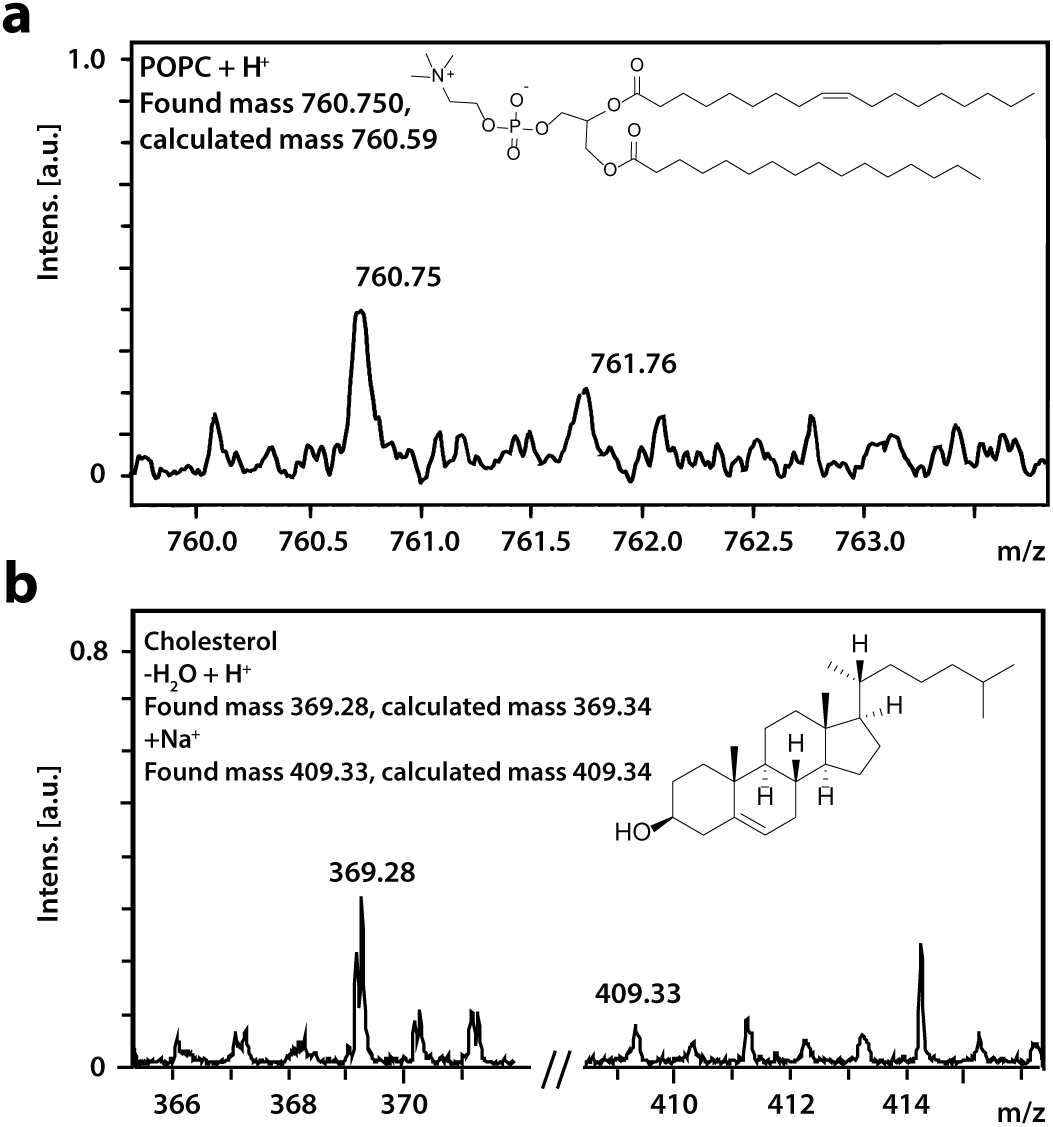
Analysis of lipids co-purified with hTRPM4 in MAASTY lipid nanodiscs by MALDI-TOF mass spectrometry. Representative spectra of identified lipids: POPC (**a**) and cholesterol (**b**).

**Figure S15:**
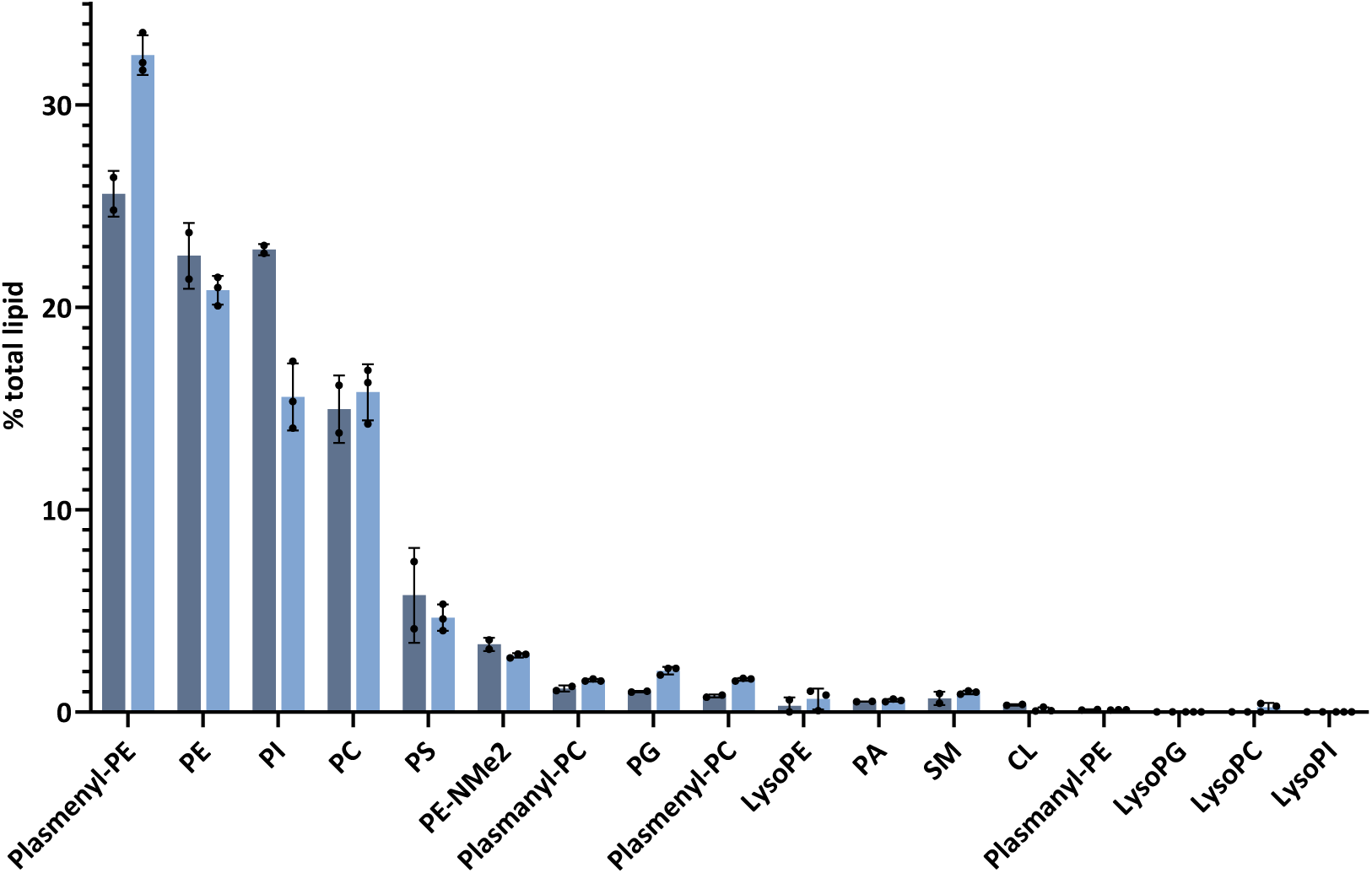
Lipidomics analysis of hTRPM4 in MAASTY lipid nanodiscs. Bar graphs show the relative abundance of phospholipids from the purified hTRPM4 lipid microenvironment analyzed from two separate protein expressions and purifications (grey and blue bars) with two and three technical replicates, respectively. Error bars represent ± SD. The phospholipids, PE, PI, PC and PS constituted the majority of observed lipid species. All the lipid types observed in the lipidomics analysis are supplied as Source Data.

**Figure S16:**
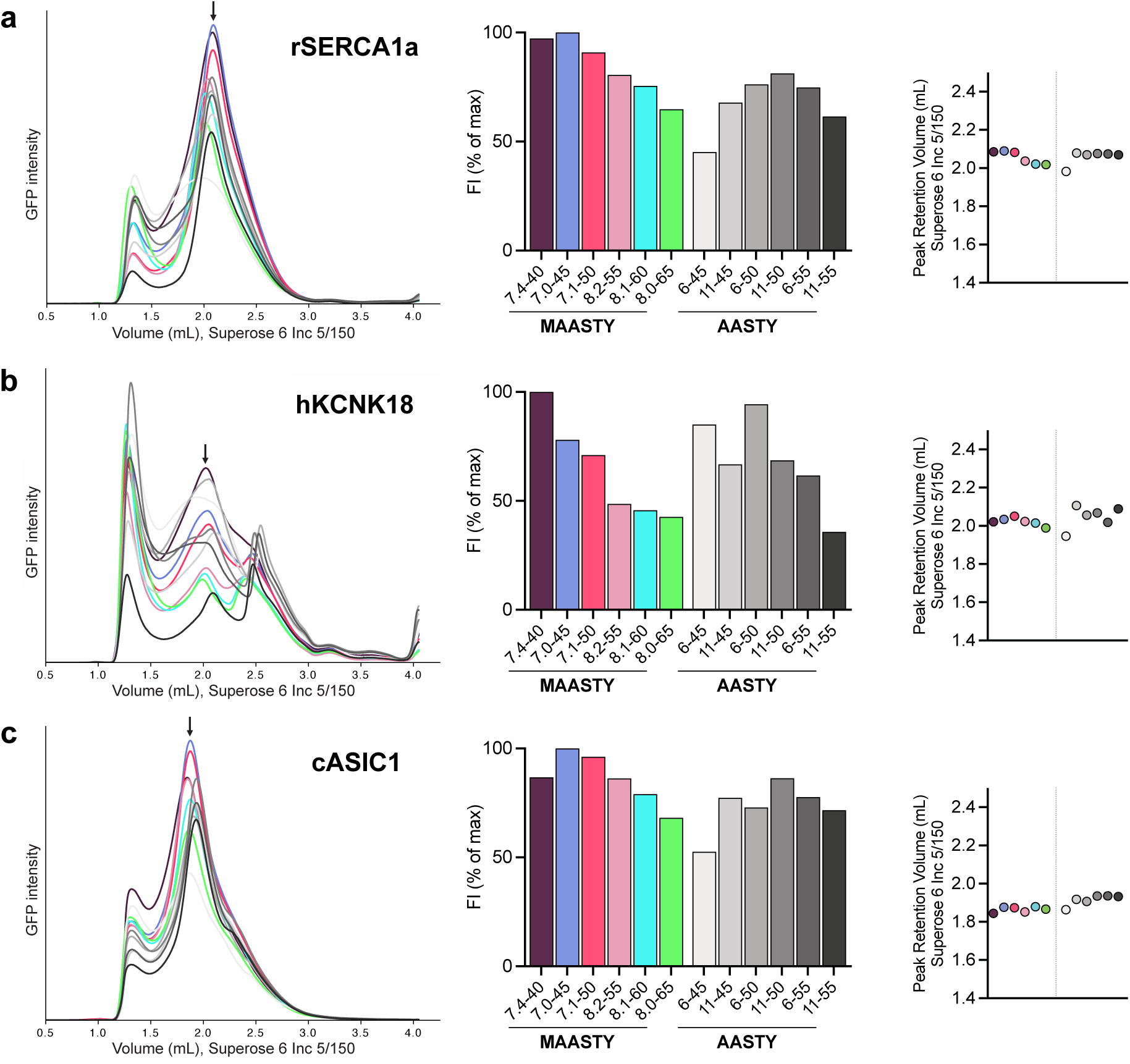
Solubilization of rSERCA1a (a), hKCNK18 (b), and cASIC1 (c) with MAASTY and AASTY copolymers, as examined by FSEC. All membrane proteins were recombinantly expressed with an eGFP fusion tag. The samples were injected onto a Superose 6 Increase 5/150 gel filtration column and and detected for eGFP with excitation at 489 nm and emission at 507 nm. The first column shows raw FSEC traces with a void peak around 1.2 mL, and arrows that denote the presumed protein-nanodisc peak. The second column displays the maximum fluorescence intensities of the denoted protein-nanodisc peak, normalized to maximum detected peak intensity. The third column exhibits peak retention volumes. Each unique membrane protein copolymer sample was injected twice or more to check reproducibility.

**Figure S17:**
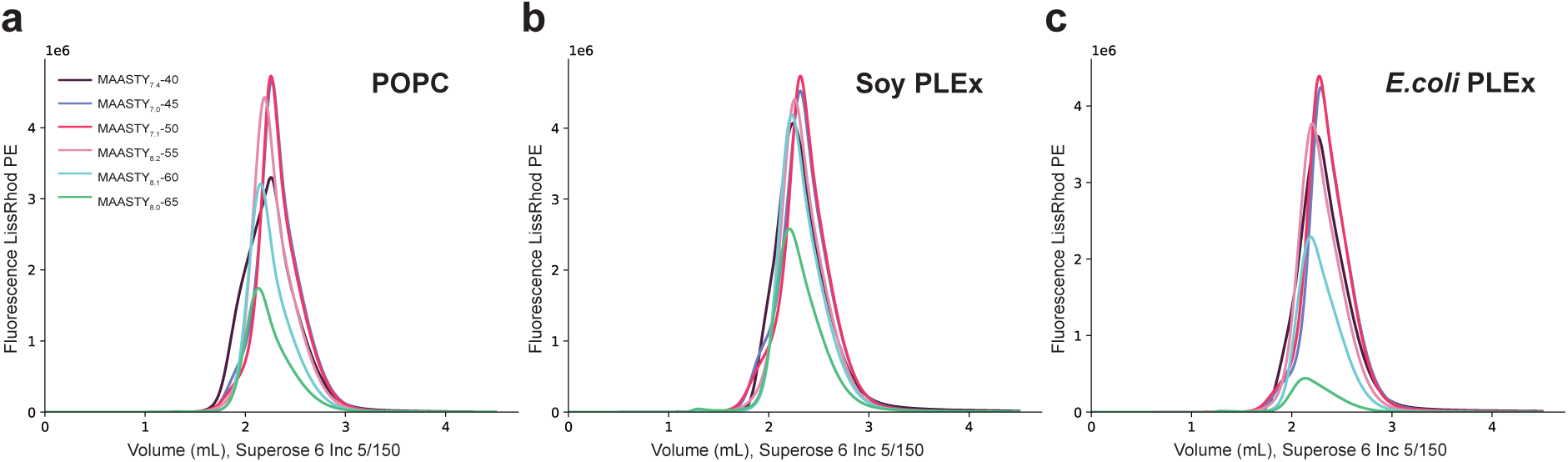
Solubilization of lipids into nanodiscs by MAASTY copolymers. FSEC traces of MAASTY-stabilized empty nanodiscs with the following lipids: POPC (**a**), Soy PLEx (**b**), and *E. coli* PLEx (**c**), mixed with 2% LissRhod PE with excitation at 554 nm and emission at 576 nm.

**Supplementary Table 1:** Data acquisition and processing, modelling statistics and validation scores.

|  | **hTRPM4 in**<br>**MAASTY** | **hTRPM4 in**<br>**MSP2N2** |
| --- | --- | --- |
| **Data collection** |  |  |
| Pixel size (Å) | 0.749/0.725 | 0.725 |
| Dose (e<sup>-</sup> Å<sup>2</sup>) | 52 | 52 |
| Defocus range (µm) | 0.5-2.5 | 0.5-2.5 |
| Total micrographs | 6308 | 7945 |
| **Data processing** |  |  |
| Curated micrographs | 6070 | 7627 |
| Picked particles | 1,306,897 | 1,104,004 |
| Selected particles (post 2D) | 126,417 | 255,586 |
| Selected particles (post 3D) | 87,697 | 103,254 |
| Map resolution (Å) | 3.46 | 3.78 |
| Map sharpening B factor (Å<sup>2</sup>) | -92 | -147 |
| **Model refinement** |  |  |
| Protein residues | 558-711, 767-835,<br>850-1089, 1107-1108,<br>1116-1166 | 558-711, 767-835,<br>842-1094, 1107-1181 |
| Ligands (pr. chain) | 3 CLR, 1 Ca<sup>2+</sup> | 3 CHS, 1 Ca<sup>2+</sup> |
| **R.M.S. deviations** |  |  |
| Length (Å) | 0.002 | 0.002 |
| Angles (°) | 0.531 | 0.618 |
| **Ramachandran** |  |  |
| Favored (%) | 96.21 | 93.30 |
| Allowed (%) | 3.79 | 6.70 |
| Outliers (%) | 0.00 | 0.00 |
| **Model validation** |  |  |
| MolProbity score | 1.70 | 2.37 |
| Clash score | 8.22 | 11.06 |
| Rotamer outliers (%) | 0.57 | 3.15 |
| Rama-Z score | 1.23 | 0.01 |

**Supplementary Table 2:** Overview of AASTY copolymers used in this study. Abbreviations: acrylic acid, AA; number-average molecular weight, M_n_.

| Nomenclature | Initial AA (%) | M<sub>n</sub> (kDa) |
| --- | --- | --- |
| AASTY<sub>6</sub>-45 | 45 | 6 |
| AASTY<sub>11</sub>-45 | 45 | 11 |
| AASTY<sub>6</sub>-50 | 50 | 6 |
| AASTY<sub>11</sub>-50 | 50 | 11 |
| AASTY<sub>6</sub>-55 | 55 | 6 |
| AASTY<sub>11</sub>-55 | 55 | 11 |

